# The cause-effect conundrum of local-scale site and soil factors in acute oak decline (AOD)

**DOI:** 10.1101/2025.01.24.634765

**Authors:** Liz J. Shaw, Mojgan Rabiey, Mateo S J Garcia, Toni Clarke, Alice Broome, Luci Corbett, Oliver R. Booth, Glyn A. Barrett, Gail M. Preston, Nadia Barsoum, Karsten Schönrogge, Robert W. Jackson, Duncan Ray

**Author notes:** These authors made equal contributions (Mojgan Rabiey, Mateo S J Garcia).

## Abstract

**Background and aims:** Acute oak decline (AOD), a decline syndrome affecting mature oaks, involves bacterial pathogens which likely act as opportunists under host stress. Trees displaying symptoms (bleeding cankers) appear in localized clusters, not whole stands. This study investigates the potential involvement of local-scale factors, in interaction with large-scale environmental drivers, in influencing onset and progression of AOD.

**Methods:** AOD-symptomatic (n=30) and asymptomatic trees (n=30) across three UK oak woodlands were assessed for tree characteristics, their surrounding context, and soil properties.

**Results:** Tree health status was linked to significant differences in soil and tree properties across sites. Symptomatic trees exhibited greater loss of crown density, lower local stand (0-20 m) basal area and shallower depth to gleying. Significant differences in soil properties included lower concentrations of Olsen P, total N, and exchangeable Mg in symptomatic trees, alongside higher exchangeable Fe, especially at 40–50 cm depth. Depth to gleying and exchangeable Fe were identified as the most influential predictors of AOD.

**Conclusions:** AOD symptomatic trees may experience seasonal soil water saturation closer to the surface compared to asymptomatic trees, resulting in a higher proportion of their roots being exposed to an anoxic, iron-reducing environment. This study is the first to report such an association between gleying depth, likely seasonal water saturation, and symptom status for AOD. It is unclear whether water balance and associated soil nutrient variations are predisposing factors or consequences of declining tree health, though the identified local-scale factors likely contribute to AOD. A feedback loop is conceptualised where declining tree health worsens soil conditions, creating a negative cycle that accelerates tree decline.

## 1. Introduction

Acute oak decline (AOD) is a decline syndrome within the wider oak decline complex affecting the native oak species (*Quercus robur* and *Quercus petraea*) in Britain that was first described by (Denman & Webber, 2009). The decline is an outcome thought to be caused by multiple contributing factors, the main visible symptom being the presence of bleeding cankers exuding from cracks in the outer bark of the trunk (Denman *et al*., 2014; Gosling *et al*., 2024). The symptoms are mainly observed on mature trees (>50 years old). Other observed tree changes include necrotic lesions in the inner bark around the seepage point and often the presence of larval galleries of *Agrilus biguttatus*, a bark beetle, but the role of this beetle in AOD has yet to be fully resolved (Reed *et al*., 2018; Cambon *et al*., 2024; Tkaczyk & Sikora, 2024). Symptoms have been shown to affect individual trees or localised clusters rather than whole stands and are associated with high tree mortality over a relatively rapid time period (3-8 years) (Brown *et al*., 2016; Denman *et al*., 2022). Some trees, however, may callus over infections and enter remission (Brown *et al*., 2016). Geographically, in the UK, AOD has mainly been reported in the Midlands and South-East England (Brown *et al*., 2017) but, since its first description in Britain, AOD has been reported more widely across continental Europe as well as Iran (Moradi-Amirabad *et al*., 2019; Gonzalez & Ciordia, 2020; González & Ciordia, 2020; Ruffner *et al*., 2020; Zalkalns & Celma, 2021; Crampton *et al*., 2022; Fernandes *et al*., 2022).

Various studies have shown that particular bacterial species within the family Enterobacteriacae, including *Brenneria goodwinii* (Denman *et al*., 2012), *Gibbsiella quercinecans* (Brady *et al*., 2010), *Rahnella victoriana* and *Rahnella variigena* (Brady *et al*., 2014) can be consistently isolated from the bleeding cankers in the stem (Denman *et al*., 2016; Denman *et al*., 2018) or are highly represented within the lesion microbiome (Sapp *et al*., 2016; Denman *et al*., 2018). Log/tree inoculation tests (Denman *et al*., 2018) showed that *B. goodwinii* and *G. quercinecans* alone can cause larger lesions than control tests, but that dual inoculations cause the largest lesions, suggesting the production of lesions is the result of a polymicrobial infection complex. Intriguingly, disease symptoms were only ever observed in up to 65% of inoculated logs/trees, suggesting that either other components of disease are required or that some variation in the tree genotype, tree condition, phenotype or microbiome can affect the development of lesions. It follows that these bacteria might be endophytes that become opportunistic pathogens. Consistent with this, *B. goodwinii* and *G. quercinecans* are not only consistently prevalent in the microbiome of canker lesions but have also been isolated and detected, albeit at low frequency (0.01% of sequence reads), in metagenome analyses of healthy oak trees (Denman et al., 2018). Subsequent incidental genetic sampling additionally suggests that AOD-associated bacteria may be members of the normal oak microbiome (Gathercole et al., 2021).

Genome analysis of these and other bleeding canker-associated bacteria (*R. victoriana, R. variigena* and also *Lonsdalea Britannica*) reveals all to have potential virulence genes to support colonisation of the plant and contribute to disease (Brady *et al*., 2012; Denman *et al*., 2018; Doonan *et al*., 2019). Whilst *B. goodwinii* has been implicated as the primary pathogen component, based on its frequency in AOD-positive lesions and having more candidate virulence factors than the other bacteria (Doonan *et al*., 2019), it is suggested that within the complexity of the canker lesion, the multiple bacterial species interact and cause disease, dependent on host condition, without an apparent primary pathogen (Broberg *et al*., 2018; Doonan *et al*., 2019). Members of the bacterial family *Enterobacteriaceae*, to which the AOD-associated bacteria belong, are known to establish a range of ecological relationships with host plants, from mutualism to opportunistic pathogenesis (Hardoim *et al*., 2015). Notably, species of the genera *Brenneria* (Maes *et al*., 2002; Maes *et al*., 2009) and *Rahnella* (Kandel *et al*., 2017) have been reported as endophytes on non-woody (Rosenblueth & Martinez-Romero, 2006; Torres *et al*., 2008) and woody (Rosenblueth & Martinez-Romero, 2006; Torres *et al*., 2008; Maes *et al*., 2009; Shen & Fulthorpe, 2015; Kandel *et al*., 2017) plant species without always causing disease symptoms. For instance, *Brenneria salicis*, typically associated with watermark disease in willow, has been found as a general endophyte in symptomless willow, poplar, and alder trees, indicating that the presence of *Brenneria* in willow does not inevitably lead to disease (Maes *et al*., 2009).

The discovery of AOD-bacteria associated with symptomless oaks and the presence of closely related bacterial strains as endophytes in other symptomless plants have prompted further discussion regarding the implications of these findings for understanding AOD etiology (Maddock *et al*., 2023). The possibility that *G. quercinecans, B. goodwinii* and *R. victoriana* are widespread endophytes that are opportunists has been previously discussed in the context of AOD and the notion that they are opportunists exploiting necrotic tissue initiated by another organism has been discounted (Denman *et al*., 2018). However, a hypothesis emerging from recent data (Doonan *et al*., 2019) suggests that these bacteria exist as endophytes within oak trees and could multiply as primary necrotic agents when trees become immunocompromised, resulting in susceptibility. In this scenario, the endophytes transition into opportunistic pathogens. If this hypothesis is correct, it raises critical questions about how and why oak trees become susceptible to these bacteria.

A general model in tree health is that decline diseases are a function of opportunistic pathogens acting on stress-weakened trees (Houston, 1992; Wargo, 1996). The model implies that changes in an ecosystem create a threshold whereby a stressed organism changes from a healthy state to a vulnerable one that can then be exploited by an opportunistic infection. Accepting the preceding argument that the canker pathogens associated with trees experiencing AOD might be widely present in oak and cause disease depending on host condition, it is likely that AOD fits this decline disease model, which involves an initial role of predisposition and inciting factors in weakening host resilience (Manion & Lachance, 1992; Denman *et al*., 2022).

A spatial study in the UK has shown AOD occurrence to be correlated with climate (rainfall, temperature) and atmospheric deposition (oxidised N, cations and S) factors, which might act in predisposition (Brown *et al*., 2018). A review of wider oak decline in Europe (Thomas, 2008) has highlighted weather extremes (drought, winter and spring frost, severe waterlogging) and defoliation by lepidopteran larval outbreaks (sometimes in combination with late powdery mildew infections), often acting in the same year or in temporal sequence, to be important inciting stressors in oak decline. Drought compromises tree condition through limiting carbon assimilation and therefore carbohydrate supply as a result of reductions in stomatal conductance that are elicited to prevent extreme water-loss (Cochard *et al*., 1996). Insect defoliation also impacts on carbohydrate reserves and can be severe for intense or prolonged outbreaks (Thomas *et al*., 2002). Similarly, spring frost, through damage to expanding leaves, depletes carbohydrate reserves and can also impact water transport through freeze and subsequent thaw of xylem in earlywood vessels (Thomas *et al*., 2002). The depletion in stored carbon reserves or hydraulic compromise could clearly undermine a tree’s ability to resist pathogen attack. However, given that individual trees within stands or localised clusters rather than whole stands (Brown *et al*., 2016) experience AOD, but weather-related factors and defoliating insect outbreaks act at population scales or larger, it is not possible to explain the distribution of symptomatic trees solely as a function of these large-scale processes.

Among-tree genotypic variation in the immune system used to either resist canker pathogens or to tolerate combinations of large-scale predisposing and inciting factors might be one explanation for the apparent mixed susceptibility of oak trees in the same woodland to acute decline. An alternative (non-mutually-exclusive) explanation is that local site factors, at the scale of the individual trees, in combination with some or all of the larger scale (e.g. climate- or weather-related) processes, create the threshold that renders certain trees susceptible to opportunistic infection by the canker bacteria. Forest soils and environments are notoriously variable in soil physical and (bio)chemical properties over scales (cm to m) that are relevant to individual trees (Farley & Fitter, 1999; Baldrian, 2014; Stursova *et al*., 2016). Local scale spatial variability arises, in part, from historical disturbances, such as wind disturbance and felling, deadwood decomposition and also variable micro-topographical environments (e.g. hollows and ridges) with distinct microclimates, soil depths, nutrient availabilities and understory species compositions and plant-soil feedbacks (Beatty, 1984; Stursova *et al*., 2016).

Local soil conditions that have been discussed as contributing to oak decline (but not specifically AOD) mainly relate to the role of soil physical properties (texture, compaction) and their interaction with weather extremes (drought or rainfall excess) in controlling the availability of water to roots (Thomas & Hartmann, 1998; Vincke & Delvaux, 2005). In addition, local properties related to the context of individual trees with respect to stand density and social status, in interaction with soil moisture status, have been shown to have complex effects on drought resilience of *Q. petraea* (Trouve *et al*., 2017). In the case of soil chemical properties, whilst we have a sound understanding of soil nutrient imbalances that might be damaging to root and tree growth in oak either directly (e.g., excess Mn^2+^ (Thomas & Sprenger, 2008) or N (Smithwick *et al*., 2013) or low Ca: Al molar ratios (Vanguelova *et al*., 2007)) or through influences on beneficial or detrimental root-associated microorganisms (Jonsson *et al*., 2005; van der Linde *et al*., 2018), the relationship between local scale soil chemical properties and oak decline has seldom been studied.

Previous attempts to relate soil chemical status to oak (*Q. petraea* and *Q. robur*) damage have focussed at the stand rather than individual tree level (Thomas & Buttner, 1998; Jonsson *et al*., 2005) and did not identify a clear impact of soil chemistry on oak health status. We are aware of only one study to date (Rozas & Sampedro, 2013) to investigate a possible link between the tree scale availability of soil nutrients and oak (*Q. robur*) decline: lower nutrient (N, Ca, Mg and Na) concentrations were found in soils surrounding dead trees when compared to those designated either ‘healthy’ or ‘declining’. In combination with tree-ring data, it was concluded that trees with lower nutrient (particularly Ca) availability had a reduced phenotypic plasticity to stress caused by water excess and therefore were more predisposed to die. Whilst this study (conducted in Atlantic wet forests in SW Spain) did not focus on trees showing the main diagnostic external symptom of AOD (i.e. bleeding cankers) but on trees classified as declining on the basis of dieback, crown transparency, epicormic shoots and leaf discolouration, it reveals the interesting link between the local availability of soil nutrients and the biological response of trees to other stress factors.

When considering the influence of local environmental conditions on oak health, it should also be considered that trees, depending on their health status, can significantly influence their surrounding environment, particularly local soil conditions (Zinke, 1962; Miles, 1986; Flinn & Marks, 2007), via altered litter inputs, nutrient uptake and water dynamics. This can complicate the task of disentangling cause from effect when analysing factors associated with AOD symptomatology. Therefore, with a focus on three forest sites in the UK, the objective of this study was to analyse soil, tree, and stand context factors at the individual tree level, with the aim of understanding their relationship with AOD symptoms while also considering potential feedback loops between tree health and soil properties. Our findings reveal significant correlations between AOD symptoms (cankers) and specific soil and tree factors, offering novel insights into the complex interactions that may drive oak decline.

## 2. Materials and methods

### 2.1 Study sites

Three oak woodland sites were selected in the south of England for detailed site investigations. At the sites there had been documented (Booth, 2021) or anecdotal reports of oak decline, featuring stems with dark necrotic bleeding cankers and dieback of the branches in the crown. The three sites, previously investigated in a study of root ectomycorrhizal communities of AOD symptomatic and asymptomatic trees (Barsoum *et al*., 2021), were: Writtle Forest (WF), Monks Wood (MW) and Stratfield Brake (SB) (Table 1). Each was typical of oak woodland in south England, at elevations below 80m, comprised of mature pedunculate oak (*Quercus robur* L.) and sessile oak (*Quercus petraea* (Mattuschka Liebl.)) in mixture with other species on clayey textured soils derived from glacial and riverine deposits. In all three woodlands, oak was the dominant species forming over 60% of the canopy cover.

**Table 1.** General characteristics of the climate and soils at the three study sites.

|  | Monks Wood (MW) | Stratfield Brake (SB) | Writtle Forest (WF) |
| --- | --- | --- | --- |
| Latitude/Longitude (degrees) | 52.4028/-0.2384 | 51.8035/-1.2815 | 51.6944/0.3718 |
| Elevation (m) | 10-20 | 50-70 | 40-60 |
| Annual precipitation (mm) 30-year average 1991-2020 | 558 (Cambridge) | 681 (Oxford) | 615 (Heathrow) |
| Mean annual temp (°C) 30-year average 1991-2020 | 10.7(Cambridge) | 11.0 (Oxford) | 11.5 (Heathrow) |
| Soil association | Evesham (411c)/Cannamore (513) | Wickham (711f) | Windsor (712c)/Essendon (714d) |
| Major soil sub-group | Clayey textured surface-water gleys | Clayey textured surface-water gleys | Clayey textured surface-water gleys |
| Hydrology of soil types (HOST) class | Slowly permeable soils with slight seasonal waterlogging and low storage capacity over impermeable clay substrates with no storage capacity (23) | Slowly permeable, seasonally waterlogged soils over slowly permeable substrate with no storage capacity (25) | Slowly permeable, seasonally waterlogged soils over slowly permeable substrate with no or negligible storage capacity (24/25) |
| Soil swelling shrinkage potential | High (4) | High (4) | High/Very high (4/5) |
| Lithology | Till, compact head, and massive clays (Jurassic & Cretaceous) | Very soft massive clays (Drift over Jurassic and Cretaceous clay or mudstone) | Till, compact head, and very soft massive clays (Plateau and river terrace gravel & Tertiary clay) |

Writtle Forest is an ancient woodland that was a medieval royal hunting forest with records from the 12^th^ Century; the woodland is comprised mainly of oak and hornbeam (*Carpinus betulus* L.). Monks Wood is an ancient woodland of oak, ash (*Fraxinus excelsior* L.) and field maple (*Acer campestre* L.) with an understorey of hazel (*Corylus avallana* L.), blackthorn (*Prunus spinosa* L.), dogwood (*Cornus sanguinea* L.), wild service tree (*Sorbus torminalis* L. (Crantz)) and willow *(Salix cinerea* L.) (Steele & Welch, 1973). Stratfield Brake is a mature secondary woodland comprised mainly of oak, with minor components of ash, field maple, hazel, hawthorn (*Crataegus monogyna* Jacq.) and elm (*Ulmus procera* Salisb.). All three sites have a similar climate, with Stratfield Brake further west receiving approximately 10% more precipitation than Writtle Forest (measured at Heathrow) and Monks Wood receiving approximately 10% less than Writtle Forest (Table 1). The mean annual temperatures are all similar. Each of the woodland sites occurs on clayey textured soils highly prone to shrinkage (in the summer) and swelling (in the winter). This causes soils to be impermeable in the wetter winter period and more permeable due to cracking in the drier summer months. Key surface soil (5-15 cm) properties for each site (organic matter, pH, texture, total C and N) are reported in Table 2 with full data for surface and deeper (40-50 cm) soil properties given in Supplementary Tables S1 and S2, respectively.

**Table 2.** Key soil physicochemical properties at the three study sites. Data are overall mean (± SD) based on soil samples taken from a depth of 5-15 cm around 20 sampled trees (10 symptomatic, 10 non-symptomatic) per site. Means for each soil characteristic sharing a letter in common are not significantly different (p<0.05) among sites, according to Games-Howell Pairwise Comparisons.

|  | Monks Wood (MW) | Stratfield Brake (SB) | Writtle Forest (WF) |
| --- | --- | --- | --- |
| Organic matter (%) | 13.5 ± 2.7<sup>A</sup> | 15.3 ± 3.1<sup>A</sup> | 16.3 ± 6.4<sup>A</sup> |
| pH (H<sub>2</sub>O) | 4.7 ± 0.7<sup>A</sup> | 3.6 ± 0.2<sup>B</sup> | 3.6 ± 0.3<sup>B</sup> |
| Clay (%) | 1.6 ± 0.8<sup>A</sup> | 0.6 ± 0.3<sup>B</sup> | 0.4 ± 0.5<sup>B</sup> |
| Silt (%) | 68.7 ± 7.8<sup>A</sup> | 56.2 ± 5.3<sup>B</sup> | 47.6 ± 10.1<sup>C</sup> |
| Sand (%) | 30.0 ± 7.9<sup>C</sup> | 43.1 ± 5.6<sup>B</sup> | 52.0 ± 10.7<sup>A</sup> |
| Total C (%) | 6.0 ± 1.5<sup>A</sup> | 7.6 ± 2.2<sup>A</sup> | 8.3 ± 4.0<sup>A</sup> |
| Total N (%) | 0.47 ± 0.1<sup>A</sup> | 0.52 ± 0.1<sup>A</sup> | 0.42 ± 0.2<sup>A</sup> |

### 2.2 In-field survey, sampling and analysis for selected study trees

#### 2.2.1 Selection of trees

The survey and sampling process was conducted July to November 2016 and targeted 10 trees with bleeding canker symptoms (hereafter, regarded as AOD-symptomatic) and 10 AOD-asymptomatic trees at each woodland site. At each site, AOD-symptomatic trees were identified through a walking survey of the area. Trees exhibiting signs of bleeding canker lesions on the lower 5 meters (as estimated by eye) of the stem were recorded, with their coordinates documented to create a comprehensive sampling frame of symptomatic trees. From this list, 10 trees were randomly selected for the study. To select asymptomatic trees (those without visible bleeding canker lesions), a random number between 1 and 60 was used to determine a direction (bearing) from each symptomatic tree. Following this bearing, the first asymptomatic tree encountered after walking a distance of >20 meters from the AOD tree was chosen. Latitude/longitude coordinates for the selected study trees is given in Supplementary Table S3.

These trees formed the focus for in-field assessment of tree and tree context properties (section 2.2.2), soil sampling for subsequent laboratory analysis of soil physical and chemical properties (section 2.2.3) and leaf sampling for laboratory assessment of leaf morphometry (section 2.2.4).

#### 2.2.2 In field tree and tree context properties

After selecting and marking symptomatic and asymptomatic trees (section 2.2.1), a survey was conducted to assess the trees and their surronding context. Standard forest mensuration procedures (Hamilton, 1975) and standard forest crown condition monitoring protocols (Lakatos *et al*., 2014) were used to measure dimensions of stem diameter (at breast height, 1.35 m), tree height, crown dimensions, the proportion of the crown intersecting neighbouring crowns, and the percentage reduction in crown density. The social status (dominant, codominant, subdominant, suppressed, dying) of the study tree was recorded. A soil pit, 0.8m deep was excavated at a randomly-located distance of 5m from each study tree. Measurements of horizon depth, soil texture, stoniness (Pyatt *et al*., 2001), and presence of roots were recorded, and the soil identified using the major soil sub-group classification (Avery, 1990). Depth to gleying measurements were made in the soil pits, where a ruler was used to determine the distance from the soil surface to the uppermost signs of grey or orange mottling, as identified according to the Munsell Soil Color Chart. Photographs were taken to verify the depths of the gleyed layers. The micro-topography of the site was measured along eight radial transects at 2.5m intervals inside the 10m radius, and at 5m intervals from the 10m to 20m radius. The site slope and aspect were also measured and recorded. Finally, basal area was measured using a relascope within a 20m and a 40m zone around the tree. Supplementary Figure S1 shows a schematic of the site survey and site assessment surrounding each study tree.

#### 2.2.3 Soil sampling and physicochemical analysis

For each study tree selected (section 2.2.1), eight soil samples (40-50 g field moist) were taken from each of two depths (5-15 cm and 40-50 cm) from randomly-selected positions 1.5-2.0 m away from the stem base: 2 depths x 8 samples depth^−1^ = 16 samples tree^−1^. Soil samples were subsequently pooled within depth to produce one composite sample per depth per tree. Pooled soil samples were homogenized by sieving (<2mm) and air-dried prior to analysis of soil physical and chemical properties, as described in the following, with the exception of a sub-sample that was stored field moist at 4 °C prior to analysis of mineral N concentration. Organic matter content was determined by mass loss on ignition (LOI; 500 °C, 16 h) and soil pH determined in deionised water (soil: water, 1: 2.5 (m/v)). Soil particle size distribution was determined using a Malvern Mastersizer 3000 hydro laser granulometer: samples were dispersed in 3.3% (m/v) sodium hexametaphosphate and 0.7% (m/v) sodium carbonate and measured in blue and red light and data reported using a Fraunhofer size distribution model. Total C and N contents were determined on ground (Pulverisette 5 Planetary Mill) samples using a FLASH CN elemental analyser (ThermoFisher Scientific). For determination of exchangeable cations (Na^+^, K^+^, Ca^2+^, Mg^2+^, Fe^2+^, Mn^2+^ and Al^3+^), soil samples were repeat extracted with 0.1M BaCl_2_ and pooled extracts analysed using a Perkin Elmer 3000 ICP-OES fitted with a cross flow nebuliser (Cools & De Vos, 2016). The cation exchange capacity (µmol(+) g^−1^ soil) and Ca-to-Al molar ratio were calculated from the sum of the concentrations of all exchangeable cations or the Ca^2+^ and Al^3+^ data, respectively. Extractable phosphorus was determined by the Olsen method (Olsen *et al*., 1954) and extractable mineral (nitrate- + nitrite- and ammonium-) N determined by extraction (1 h, 20°C) with 1M KCl (soil:KCl, 1: 5 (m/v)) and subsequent colorimetric analysis of extracts (Skalar SAN++ continuous flow analyser). Techincal replication was included for the analysis as follows: LOI (3), pH (3), particle size distribution (5), Total C and N (2), exchangeable cations (2), Olsen P (2), Mineral N (2). In the case of KCl extractions for nitrate-+nitrite-N and BaCl_2_ extractions for Na^+^, Fe^2+^, Mn^2+^ and Al^3+^, a low percentage of samples recorded concentrations in the extract below the detection limit for the analysis. Where this was the case, concentrations were entered in to calculations as half the detection limit (USEPA, 2000).

#### 2.2.4 Leaf morphometry and characterization of oak species

From each study tree, five randomly located leaves were taken from the outer part of the canopy for morphometric and genetic analysis to ascertain the species of oak being studied. The morphometric analysis involved measuring the lamina, petiole, sinus, number of lobes, intercalary veins, shape of the basal auricle, and leaf pilosity. From these measurements the species, *Quercus robur*, *Quercus petraea* or a hybrid, could be made based on the method published by (Kremer *et al*., 2002). The genetic method for species designations were made using a Sequenom analysis at INRA, France and SNPs for the species identification are described in (Guichoux *et al*., 2013).

### 2.3 Statistical analysis

Statistical analysis of tree, tree context and soil physico-chemical properties (5-15 and 40-50 cm depths) was mainly carried out using the Statsmodels module in Python (Seabold & Perktold, 2010). Data were tested for normality of residuals using the Jarque-Bera test and for heteroscedasticity using the Breusch-Pagan test. Raw (non-transformed) data for many variables failed to meet either normality or equality of variances or both. We therefore performed Robust (to unequal variance) type III Two-Way ANOVA with site (WF, MW, SB) and health status (symptomatic, asymptomatic) as factors. Data were Box-Cox transformed: (Y^λ^-1)/ λ where λ was chosen so as to minimise the p-value testing normality of residuals (using Jarque-Bera). Significant differences were accepted at p<0.05. Where the Two-Way ANOVA identified significant effects, post-hoc comparisons were made using the Games-Howell Method and 95% Confidence in Minitab 20. In addition, the magnitude of the difference between two means was determined as Cohen’s d effect size: x1 – x2/s, where x1 and x2 are the means of the two groups being compared and s is the unweighted pooled standard deviation of the two groups.

The ‘gbm’ package in R was used to apply generalized boosted regression models (GBM) to tree health status (Greenwell *et al*., 2020) using all tree context and soil physico-chemical variables as predictors. GBM are a type of machine learning algorithm which can take multiple predictors to sequentially fit regression trees with the aim of minimizing a loss function. This process allows the most influential set of predictors to be identified and can allow for out-of-sample prediction. GBM were applied with a Bernoulli distribution, allowing two-way interactions between predictor variables and a minimum of two observations per node. Cross validation determined the optimum number of regression trees to be approximately 550. A training set was created by selecting 70% of the samples at random (selecting equally from health status and site) with the remainder of the samples used as a validation set to test prediction accuracy. Prediction accuracy was determined by calculating the area under the curve (AUC), sensitivity and specificity of the models using the ‘caret’ package (Kuhn, 2021). As there was no independent sample available for prediction this process was repeated 100 times and the average prediction accuracy calculated across all runs.

The top twenty variables, in terms of relative influence, were taken from each GBM and subsequently ranked according to the number of times they were selected from each GBM run (1-100). These top variables were then sequentially added to a generalised linear regression model (GLM) with a binomial distribution and a logit link function which predicted tree health status. Using forward stepwise regression, variables were taken from the ranking list and included in the model until newly included variables were not significant predictors of disease status (p< 0.05). Model fit was assessed using residual diagnostic functions (DHARMa package (Hartig, 2020)). Estimated marginal means were calculated for each of the significant variables using the ‘emmeans’ package (Lenth, 2021) and responses back transformed from the logit scale for ease of interpretation.

## 3. Results

### 3.1 Site properties

Across all three sites (Monks Wood, Stratfield Brake and Writtle Forest), surface soils (5-15 cm depth) had similar organic matter content and total C and N contents (Table 2) but differed with respect to site across other soil physico-chemical properties (Table S1, Figure 1). Soils from the Monks Wood site were less acidic (pH_H2O_ 4.7) than the other two sites (pH 3.6) and were the finest textured (silt + clay content ∼70%). In addition, Monks Wood soils had the highest Cation Exchange Capacity with highest concentrations of exchangeable basic cations (Ca, Mg, Na) but amongst the lowest exchangeable concentrations of acidic cations (Al, Fe, Mn) and Olsen P. Stratfield Brake soils were intermediate in their silt and clay content, CEC and concentration of exchangeable Ca and Na but had the highest exchangeable Aluminium (1.7- and 2.8- fold higher than Writtle Forest and Monks Wood, respectively). Writtle Forest soil contained over double the concentration of exchangeable Mn when compared to the other two sites. Soils sampled from 40-50 cm displayed similar trends in properties to surface soils with respect to site for pH, texture, CEC and exchangeable cations but with between-site differences for some properties becoming less (Mn and Olsen P) or more (total mineral N, organic matter content and total N) pronounced (Supplementary Table S2, Supplementary Figure S2).

**Figure 1.**
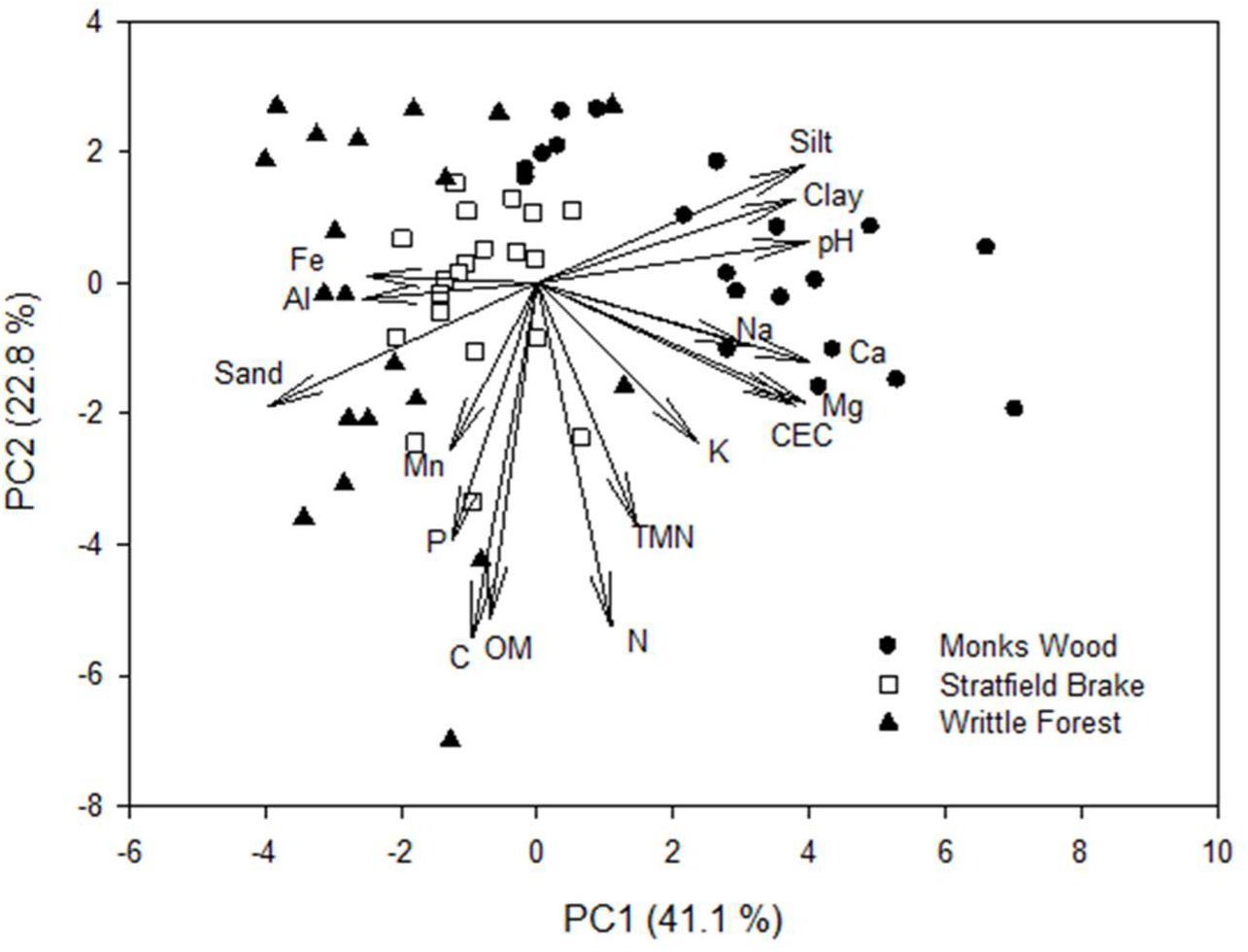
Principal Components Analysis of the study sites (symbols) and soil parameters (vectors) for surface (5-15 cm) soils. One-Way ANOVA on PC1 scores and Games-Howell pairwise comparisons revealed that all three sites differed significantly (P<0.05) from one another.

The Ecological Site Classification (ESC) (Pyatt *et al*., 2001) shows all three sites are suitable for oak woodlands (*Q. robur* and *Q. petraea*). *Quercus robur* is generally considered suitable on surface water gleys in central and southern England, that can be waterlogged during winter periods and may be slightly acid with Medium to Rich soil nutrient regime. *Quercus petraea* is not as well suited to waterlogged soils but may tolerate waterlogging for shorter periods.

### 3.2 Tree properties and AOD symptoms

Species classification based on genetic analysis of leaf samples identified that all trees from the Stratfield Brake and Monks Wood sites were *Quercus robur* (although one tree from Monks Wood did not yield DNA of sufficient quality for analysis but morphometric analysis suggested a classification of *Q. robur*) (Supplementary Table S3). For Writtle, five trees were classified as *Quercus petraea*, while the remaining 14 trees were *Q. robur* (with the species identity for one tree not determined). For the trees sampled, *Q. robur* had a relatively even distribution of symptomatic (26) and asymptomatic (28) trees. In contrast, *Q. petraea* exhibited a higher proportion of symptomatic trees (4 out of 5) (Table S3).

The oak trees under study differed between sites (p<0.05) with respect to tree height, stem diameter, rooting depth and crown density but not in crown width (Table 3). Overall, Writtle Forest trees were larger in terms of height and stem diameter and had least reduction in crown density. Across all sites, mean rooting depths were between 70 and 80 cm but with trees at Stratfield Brake with slightly (∼5-8 cm) shallower roots than at other sites (Figure 2). Examining health status as a factor revealed that trees with bleeding cankers had greater loss of crown density (p<0.001), with differences between asymptomatic and symptomatic trees being particularly distinct at Writtle Forest.

**Figure 2.**
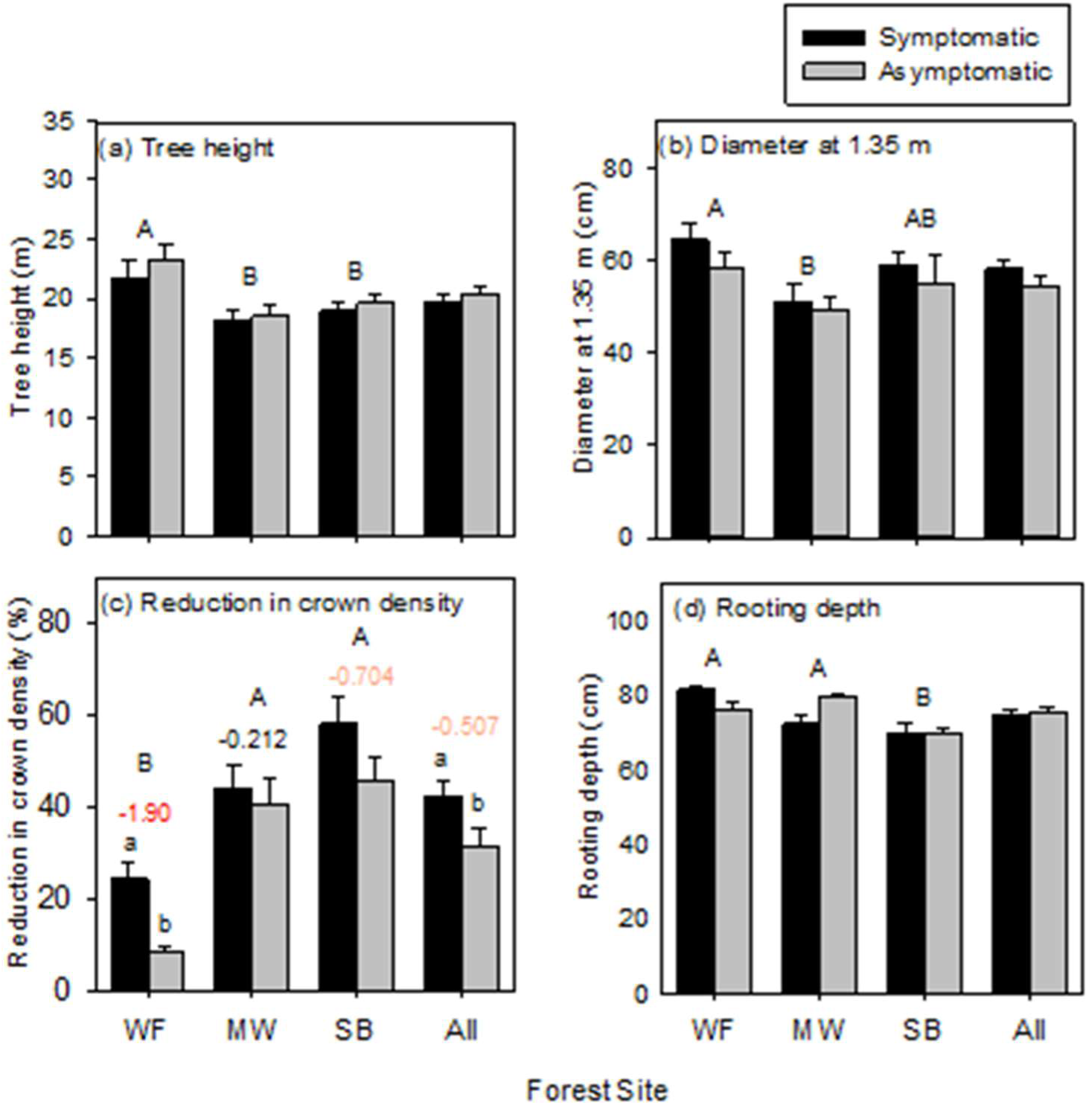
Properties of study trees (mean ± SE) that were either symptomatic (n=10) or asymptomatic (n=10) at each of the study sites: Monks Wood (MW), Writtle Forest (WF), Stratfield Brake (SB) and all three sites combined (All). Different upper case letters indicate significant differences (p<0.05) between sites whilst different lower case letters indicate significant differences between symptomatic and asymptomatic classes according to ANOVA (across all sites) or Games-Howell Pairwise comparisons (within individual sites). For reduction in crown density (c), where there was a significant effect of health status, numbers above each pairwise comparison are Cohen’s d effect size colour coded as 0.5<d<0.8: ‘medium effect’ (orange) and d>0.8: ‘large effect’ (red).

**Table 3.** Summary of outcome of Two way ANOVA* p-values examining site, tree health status and site x health status as factors explaining differences in properties of the study trees.

|  | Site (S) | Health Status (H) | S x H |
| --- | --- | --- | --- |
| *Degrees of Freedom* | 2 | 1 | 2 |
| Tree Height | **0.005** | 0.380 | 0.908 |
| Stem diameter at 1.35 m | **0.010** | 0.184 | 0.818 |
| Crown width (East-West) | 0.112 | 0.656 | 0.895 |
| Crown width (North-South) | 0.101 | 0.948 | 0.776 |
| Reduction in crown density | **<0.001** | **<0.001** | **0.002** |
| Root depth | **<0.001** | 0.749 | **0.008** |

\* ANOVA was performed in all cases with Box-Cox transformed data where the constant (lambda) was chosen to minimise the p-value testing normality of the residuals (using Jarque Bera test). Error term degrees of freedom = 54. P-values <0.05 are in bold.

### 3.3 Soil physico-chemical and tree context properties as related to AOD symptom status

#### 3.3.1 Soil physico-chemical properties as response variables to AOD symptom status

The study sites were distinct with respect to many soil physico-chemical properties measured at 5-15 cm and 40-50 cm depth (Table 2, S1, S2; Fig. 1, S2, Table 4). Examining the variation in soil properties with respect to bleeding canker symptoms revealed significant, and consistent across sites (main), effects in the case of Olsen P (5-15 cm depth), exchangeable Mg (5-15 cm and 40-50 cm), pH, organic matter content, total N and exchangeable Fe (all for 40-50 cm depth) (Table 4). In addition, pH (5-15 cm), exchangeable Al (5-15 cm and 40-50 cm), silt content (40-50 cm) and ammonium N (40-50cm) were also significantly different depending on health status but the magnitude and/or the direction of the effect was dependent on site, consistent with significant site*health status interaction terms in the ANOVA (Table 4). For cation exchange capacity at 40-50 cm depth, there was no overall association with health status, but health status was associated with this property depending on site (Table 4). Soil properties significantly predicted by oak health status as a main effect are shown in Figure 3A & B.

**Figure 3.**
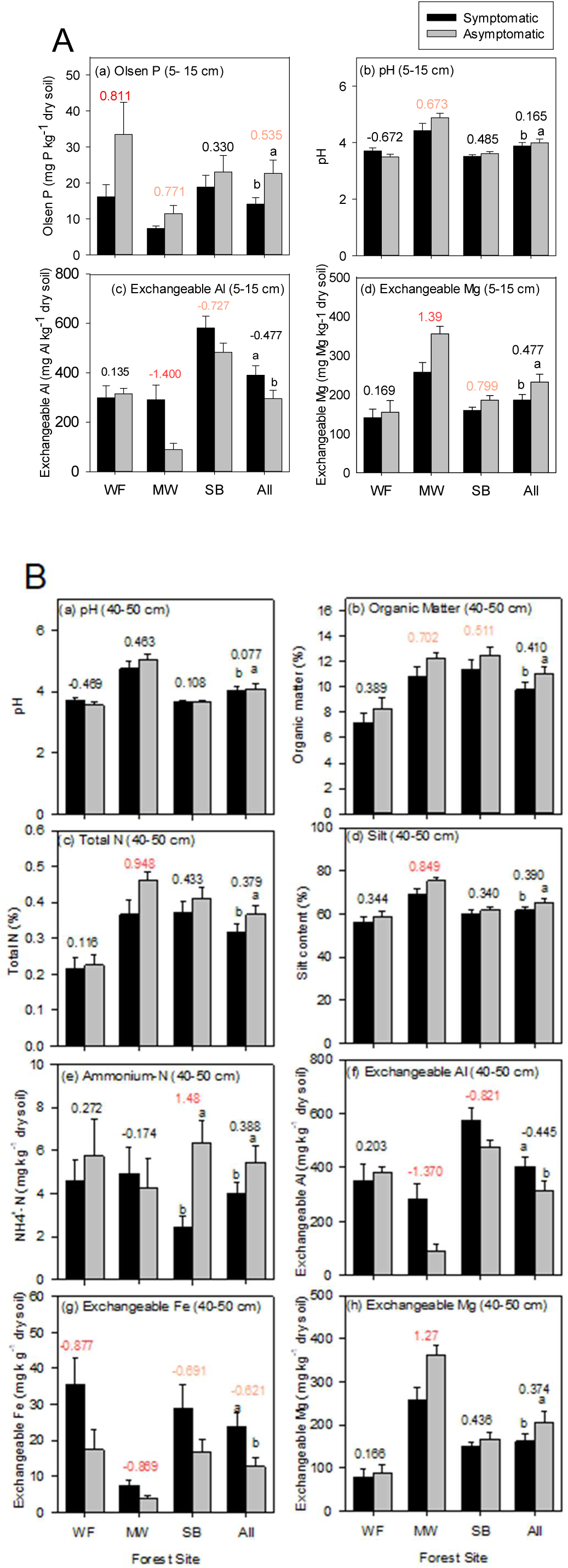
Physico-chemical characteristics (mean ± SE) of soil sampled from (A) 5-15 cm or (B) 40-50 cm) depth around trees that were either symptomatic (n=10) or asymptomatic (n=10) at each of the study sites: Monks Wood (MW), Writtle Forest (WF), Stratfield Brake (SB) and all three sites combined (All). Different letters indicate significant differences (p<0.05) between health class according to ANOVA (across all sites) or Games-Howell Pairwise comparisons (within individual sites). Numbers above each pairwise comparison are Cohen’s d effect size colour coded as 0.5<d<0.8: ‘medium effect’ (orange) and d>0.8: ‘large effect’ (red).

**Table 4.** Summary of outcome of Two way ANOVA* examining site, tree health status and site x health status as factors explaining differences in soil physico-chemical properties at each of the two soil depths studied.

|  | 5-15 cm depth |  |  | 40-50 cm depth |  |  |
| --- | --- | --- | --- | --- | --- | --- |
|  | Site (S) | Health Status (H) | S x H | Site (S) | Health status (H) | S x H |
| *Degrees of freedom* | 2 | 1 | 2 | 2 | 1 | 2 |
| Organic matter (%) | 0.128 | 0.094 | 0.735 | **<0.001** | **0.040** | 0.943 |
| pH (H<sub>2</sub>O) | **<0.001** | **0.043** | **0.034** | **<0.001** | **0.024** | 0.274 |
| Clay (%) | **<0.001** | 0.756 | 0.915 | **<0.001** | 0.753 | 0.636 |
| Silt (%) | **<0.001** | 0.956 | 0.152 | 0.171 | **0.001** | **0.002** |
| Sand (%) | **<0.001** | 0.983 | 0.141 | **<0.001** | 0.090 | 0.879 |
| Total C (%) | **0.011** | 0.158 | 0.987 | **0.039** | 0.186 | 0.799 |
| Total N (%) | 0.084 | 0.052 | 0.819 | **<0.001** | **0.039** | 0.493 |
| Nitrate-N (mg kg<sup>-1</sup>) | **0.012** | 0.497 | 0.426 | **<0.001** | 0.307 | 0.793 |
| Ammonium-N (mg kg<sup>-1</sup>) | 0.125 | 0.437 | 0.426 | 0.677 | **0.005** | **0.012** |
| Olsen P (mg kg<sup>-1</sup>) | **<0.001** | **0.029** | 0.660 | **<0.001** | 0.229 | 0.269 |
| CEC (mmol<sub>c</sub> kg<sup>-1</sup>) | **<0.001** | 0.296 | 0.080 | **<0.001** | 0.753 | **0.030** |
| Exchangeable Al (mg kg<sup>-1</sup>) | **<0.001** | **0.038** | **0.049** | **<0.001** | **0.044** | **0.045** |
| Exchangeable Ca (mg kg<sup>-1</sup>) | **<0.001** | 0.080 | 0.191 | **<0.001** | 0.079 | 0.127 |
| Exchangeable Fe (mg kg<sup>-1</sup>) | **<0.001** | 0.116 | 0.758 | **<0.001** | **0.014** | 0.596 |
| Exchangeable K (mg kg<sup>-1</sup>) | **0.003** | 0.077 | 0.319 | **<0.001** | 0.120 | 0.092 |
| Exchangeable Mg (mg kg<sup>-1</sup>) | **<0.001** | **0.004** | 0.216 | **<0.001** | **0.020** | 0.203 |
| Exchangeable Mn (mg kg<sup>-1</sup>) | **<0.001** | 0.707 | 0.372 | **0.017** | 0.481 | 0.051 |
| Exchangeable Na (mg kg<sup>-1</sup>) | **<0.001** | 0.645 | 0.324 | **<0.001** | 0.192 | 0.262 |
| Bulk density (g cm<sup>-3</sup>) | **<0.001** | 0.832 | 0.601 | nd | nd | nd |

\* ANOVA was performed in all cases with Box-Cox transformed data where the constant (lambda) was chosen to minimise the p-value testing normality of the residuals (using Jarque Bera test). Error term degrees of freedom = 54. P-values <0.05 are in bold. nd = not determined

For the properties where there was a main, and consistent across sites, effect of health status, calculation of Cohen’s *d* revealed health status to have the largest effect (*d*>0.5) on Olsen P (5-15 cm) and exchangeable Fe (40-50 cm): asymptomatic trees had 1.6 x greater mean concentrations of Olsen P, but, 0.5 x lower concentrations of exchangeable Fe across all sites. Exchangeable Mg (5-15 cm and 40-50 cm), organic matter content (40-50 cm), total N (40-50 cm) and silt content (40-50 cm), although consistent in the trend of higher mean concentrations for asymptomatic trees, were more variable in effect sizes between sites. For example, the effect size for mean exchangeable Mg was pronounced at Monks Wood (*d* >1.2) but trivial (*d* < 0.2) at Writtle Forest for both soil depths. The other properties (pH and exchangeable Al (both depths) and ammonium-N (40-50cm)) that were significantly different overall according to health status, had effects that varied in both size and sign on a site basis. For example, effect sizes for mean exchangeable Al concentration at both depths were negative (i.e. mean symptomatic > mean asymptomatic) and moderate to large (*d* >0.7) for Monks Wood and Stratfield Brake, but, trivial (*d* <0.2) and positive (i.e. mean asymptomatic > mean symptomatic) for Writtle Forest. Similarly, contrasting within-site effects for pH were recorded: soils (5-15 cm) for symptomatic trees were on average less acidic (by 0.22 pH units) at Writtle Forest but more acidic (by 0.46 pH units) at Monks Wood. The overall effect on pH, however, was small (*d*<0.2) across sites and depths with the difference in mean pH between symptomatic and asymptomatic trees being ∼0.1 pH unit or less.

Overall, symptomatic trees had less available soil P and Mg at the 5-15 cm soil depth and lower silt, total N, organic matter and Mg at 40-50 cm depth. However, concentrations of exchangeable Fe were higher for symptomatic trees (40-50 cm), as was the case for exchangeable Al (both depths) but only for two out of three sites. pH response to health status was variable and mostly subtle on the (logarithmic) pH scale.

#### 3.3.2 Tree context properties as response variables to AOD symptom status

The results for in-field-observed parameters reflecting the context of the studied trees with respect to site and AOD symptom status are shown in Figure 4. The majority of (≥ 70-%) of the trees studied were co-dominant/ dominant with the remainder classified as subdominant; the exception to this was the symptomatic sample at Writtle Forest which were drawn from a population with a greater (p=0.019; Mood’s Median Test) proportion of trees in the subdominant and suppressed classes (Figure 4a) than the asymptomatic sample. Two-way ANOVA with site and health status as factors (Table 5) revealed that symptomatic trees had a soil depth to the first signs of gleying that was significantly shallower and also a local stand basal area (measured to a distance of 20 m from focal tree) significantly smaller than for asymptomatic trees (Figures 4b &c). These effects were consistent between sites and large in size in the case of depth to gleying (*d* >1.7). In contrast, the basal area (20-40 m from focal tree) did not vary with health status (Table 5). According to the compound topographic index (CTI), trees at Monks Wood were, on average, in positions with greater water accumulation potential (mean CTI = 9.65 ± 0.84) as compared to Stratfield Brake and Writtle Forest (mean CTI = 5.19 ± 0.55 and 4.56 ± 0.80, respectively), but, the CTI was not related to tree health status (Table 5).

**Figure 4.**
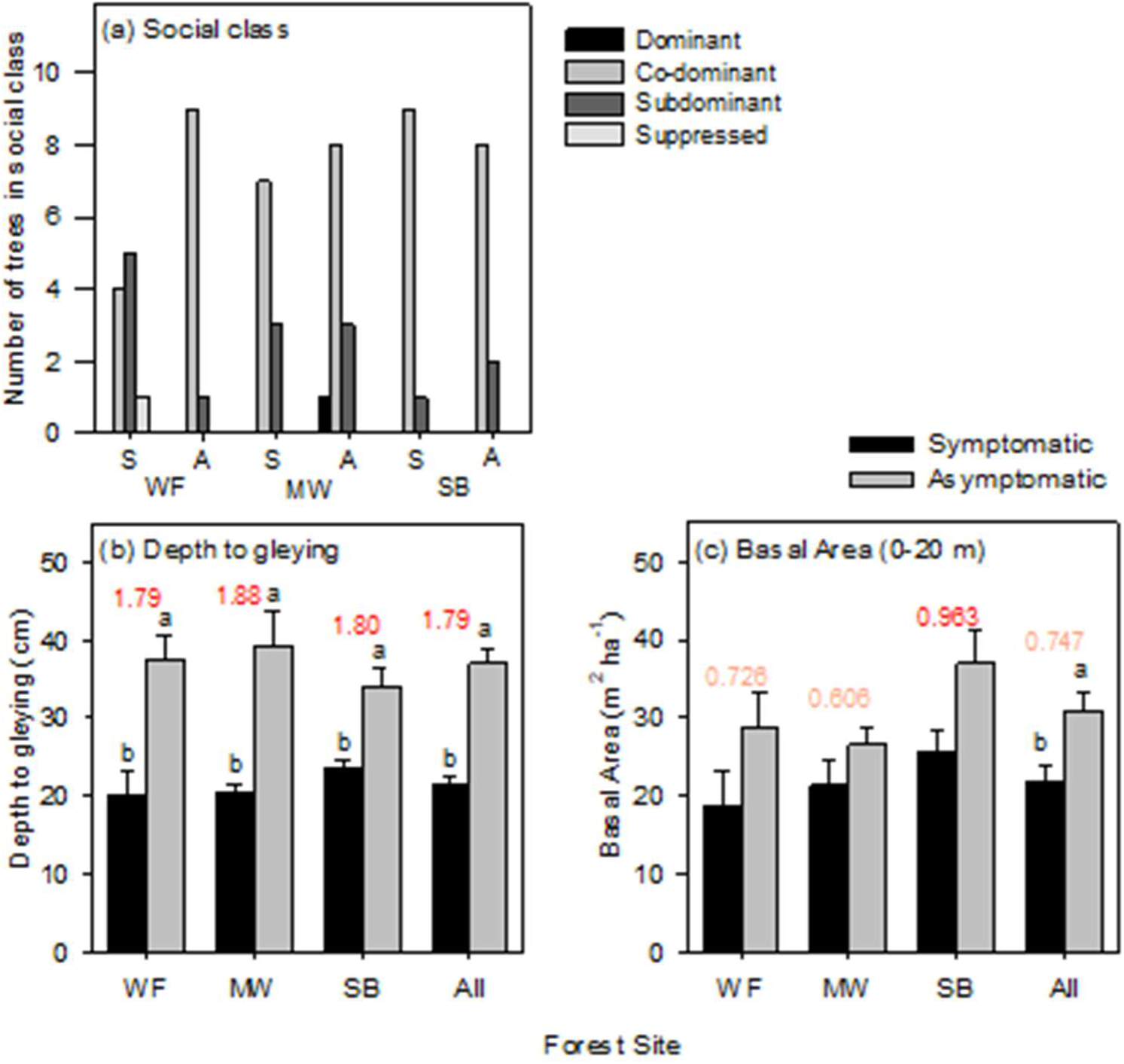
Context characteristics for trees that were either symptomatic (n=10) or asymptomatic (n=10) at each of the study sites: Monks Wood (MW), Writtle Forest (WF), Stratfield Brake (SB) and all three sites combined (All). Data for social class (a) are the number of symptomatic (S) or asymptomatic (A) trees at each site in each of four social classes. Data for depth to gleying (b) and basal area (c) are mean ± SE. Different letters indicate significant differences (p<0.05) between health class according to ANOVA (across all sites) or Games-Howell Pairwise comparisons (within individual sites). Numbers above each pairwise comparison are Cohen’s d effect size colour coded as 0.5<d<0.8: ‘medium effect’ (orange) and d>0.8: ‘large effect’ (red).

**Table 5.**
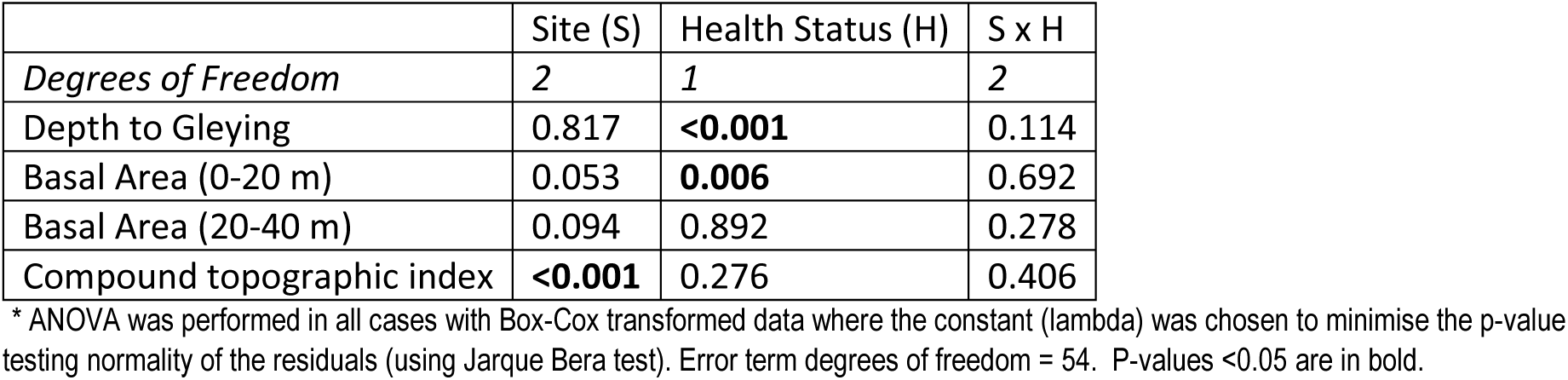
Summary of outcome of Two way ANOVA* p-values examining site, tree health status and site x health status as factors explaining differences in site properties of the study trees.

#### 3.3.3 Soil physicochemical and tree context properties as predictors of AOD symptom status

The relative influence of all soil physicochemical and tree context properties across sites was explored using GBM. Using all predictors with a non-zero influence from the GBM tree, health status was predicted in each of the 100 randomly selected validation datasets. The mean AUC for these predictions was 0.88 (s.d. = 0.06), the mean specificity 0.92 (s.d. = 0.09) and mean sensitivity 0.82 (s.d. = 0.11) (Supplementary Table S4). In other words, the sensitivity of the model indicates that, on average, 82% of diseased trees were correctly identified, while the specificity indicates that 92% of healthy trees were correctly identified.

The variables with the greatest relative influence are shown in Supplementary Table S6. Depth to gleying was amongst the top 20 predictors of all 100 GBM runs. Exchangeable Fe (40-50 cm) was selected by 95% of models, Olsen P (40-50 cm) by 90% and local stand (0-20 m) basal area by 88%. Forward stepwise regression found only depth to gleying (Anova p = 7.9 x 10^−10^) and Exchangeable Fe (40-50 cm) (Anova p=0.02) to be predictors of health status when sequentially added to the model as the addition of Olsen P (40-50 cm) was non-significant (Supplementary Table S5). Figure 5a shows the estimated proportion of diseased trees at various values of depth to gleying (cm). These values are averaged over values of Exchangeable Fe (40-50 cm), which is also present in the regression model. As the depth to gleying increases the proportion of diseased trees decreases. Figure 5b shows the estimated proportion of diseased trees at various values of Exchangeable Fe (40-50 cm), averaged over values of depth to gleying. As Exchangeable Fe (40-50 cm) values increase so does the proportion of diseased trees.

**Figure 5.**
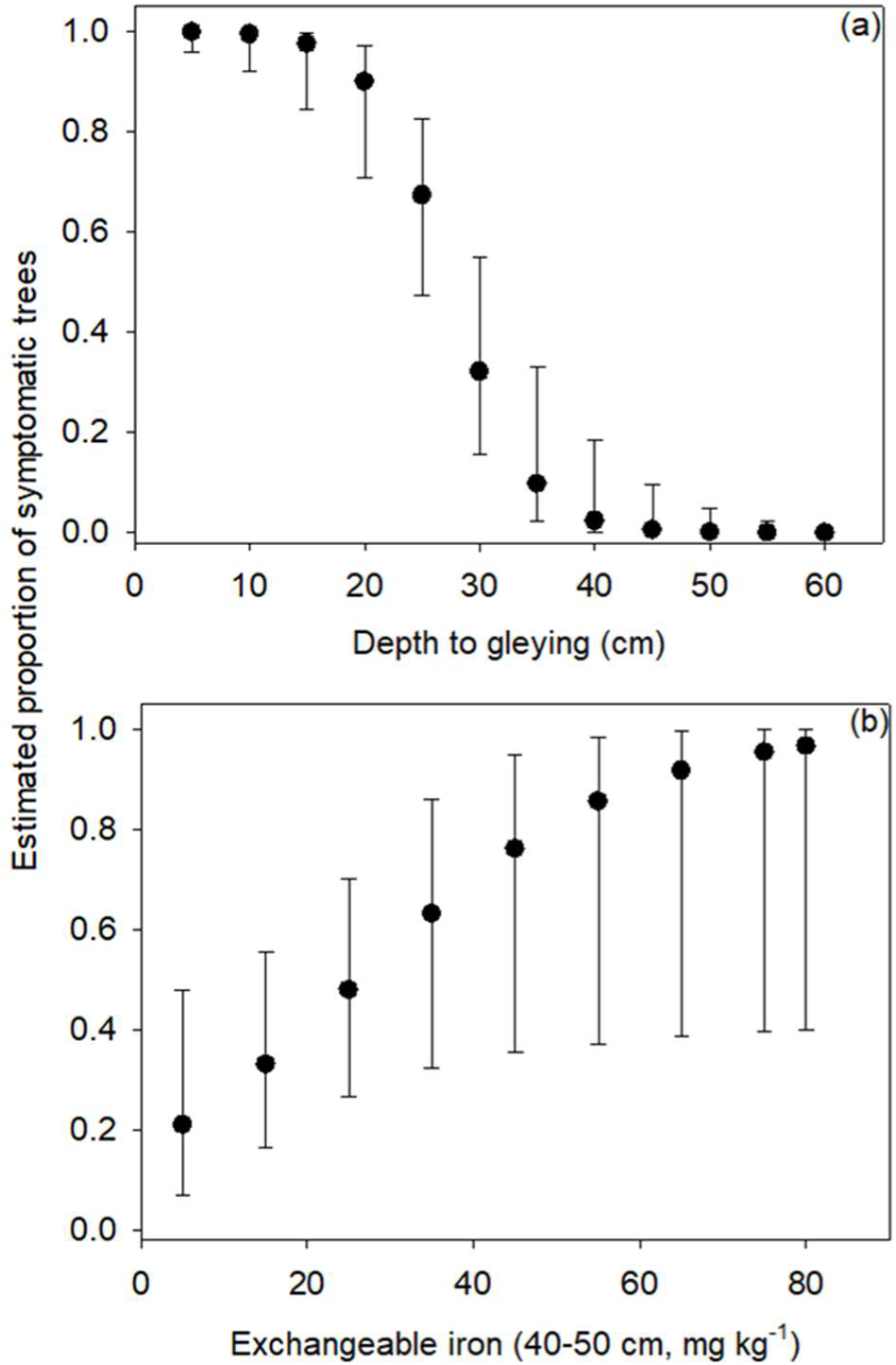
Estimated proportion of symptomatic trees at different values of (a) depth to gleying (cm) and (b) Exchangeable Fe (40-50 cm) (mg kg^−1^). Error bars represent 95% confidence intervals.

## 4. Discussion

AOD is a progressive disease in individual trees, often culminating in death. A key hypothesis is that AOD fits the classic model of a decline disease, where opportunistic pathogens exploit weakened trees. This AOD decline disease spiral is driven by multiple stressors that reduce tree vitality, making them more susceptible to bacterial infection. While research has focused primarily on the bacteria responsible for bleeding canker lesions, there are significant gaps in our understanding of other factors that contribute to this decline. AOD affects individual trees rather than entire stands. This localized impact suggests that individual tree-scale changes in the immediate environment may play a role through interaction with the tree’s physiological state to create conditions conducive to bacterial infection and decline progression. Building on this understanding, this study provides novel insight into the associations of small (individual-tree)-scale factors with AOD symptoms, specifically bleeding cankers, in native oak trees (primarily *Quercus robur*) across three representative forest sites in the UK. Across the three sites, depth to gleying and the concentration of exchangeable iron at a depth of 40-50 cm were revealed as significant factors related to tree health. After adjusting for site, AOD-symptomatic trees were also found to have reduced crown density, local stand basal area, soil organic matter concentration, and lower soil nutrient levels in the rooting zone with phosphorus, magnesium and nitrogen being notable as consistently affected across sites.

### 4.1 Tree properties as related to AOD symptom status

Our study reveals that oak trees exhibiting symptoms of AOD (bleeding cankers) tend to have lower crown density compared to asymptomatic trees. Lower crown density is widely regarded as a marker of compromised tree health, and this observation aligns with other descriptions of AOD, where symptomatic trees are often found in lower crown classes, indicating poorer health relative to the general tree population at a site. For example, research by Brown *et al*. (2016) found that AOD-symptomatic oaks are more frequently located in lower crown classes, with significant differences in crown condition across sites. Their study also reported a correlation between crown condition and AOD severity, with more frequent stem bleeds observed in trees with poorer crowns.

Reduced crown density may reflect a tree’s exposure to environmental predisposition factors and inciting stressors that weaken its defences, thereby increasing susceptibility to AOD pathogens. For instance, Kint *et al*. (2012) demonstrated that forest structure and soil fertility influence the internal stem morphology of pedunculate oaks and, by extension, their overall health. Although their focus is on stem morphology, their findings emphasize the role of environmental factors in tree health, which is consistent with our observations that crown density may also reflect underlying soil and site conditions associated with AOD symptoms.

Conversely, crown density reduction may also be a direct response to infection by the AOD bacteria. The bacteria disrupt the xylem in infected oak trees (Denman *et al*., 2022), potentially impairing water transport. If this results in stomatal closure in response to hydraulic stress, limitation of photosynthesis would reduce carbon intake. This combination of impaired water (and nutrient) transport, reduced photosynthesis, and potentially increased energy expenditure for defence or repair would restrict leaf production and lead to lower crown density.

Overall, these findings highlight the complexity of the relationship between crown density, tree health, and environmental factors that underpins the interpretation of susceptibility and progression of AOD in oak trees.

### 4.2 Soil physico-chemical and tree context properties

Both *Q. robur* and *Q. petraea* are known for their deep rooting (Rosengren *et al*., 2006). Our finding that mean rooting depths (between 70 and 80 cm) did not differ between symptomatic and asymptomatic trees, aligns with another study of oaks on hydromorphic soils (Thomas & Hartmann, 1998). In that study, fine (<2 mm diameter) and small/medium (>2 mm) roots were consistently detected at depths of 70-100 cm for both healthy and declining trees, as assessed by crown damage. Given these rooting depths, differences detected in the physicochemical properties of soil sampled at 5-15 cm and 40-50 cm depth, as well as the observations of gleying around 20-40 cm deep, fall well within the oak rooting zone. This implies that soil conditions at these depths could potentially influence, and be influenced by, the oak roots.

#### 4.2.1 Associations with oak health

Gleying, a pedogenic process caused by prolonged soil waterlogging (Brady & Weil, 2013), was identified as a significant factor in the differences between symptomatic and asymptomatic trees. This process involves the microbial reduction of Fe(III) to Fe(II) (Brady & Weil, 2013), resulting in characteristic soil mottling and grey colouration. The detection of these signs of gleying at depths shallower than 40 cm at our sites (Fig. 4) is consistent with the classification of the soils as surface-water gleys (Avery, 1980) which rest on slowly permeable or impervious clay (Table 1), making likely the formation of a perched water table in Autumn through to Spring.

The shallower depth to gleying for symptomatic trees thus suggests that these trees, on a seasonal basis, experience conditions of soil water saturation that are closer to the surface than asymptomatic trees and thus have a greater proportion of their root volume impacted by an anoxic iron-reducing soil environment. This is important because more than 60% of the cumulative root distribution for oak is typically found below 20 cm, with 40% below 40 cm (Rosengren *et al*., 2006)— the approximate depths at which gleying was observed in symptomatic and asymptomatic trees, respectively. Given that gleying converts solid-phase Fe(III) oxyhydroxides into water-soluble and exchangeable Fe(II), the significantly elevated concentrations of exchangeable iron at 40-50 cm for symptomatic trees are likely a direct result of this redox process (Gotoh & Patrick Jr., 1974). This study is the first to report an association between shallower gleying depth, likely seasonal water saturation, and symptom status for AOD specifically. However, our results align with other studies on hydromorphic soils showing that oak trees in decline (i.e. marked crown defoliation and twig abscission) are found in soils with shallower water stagnation depths (Thomas & Hartmann, 1996; Thomas & Hartmann, 1998)

In contrast to the increased exchangeable iron, the availability of P and Mg was reduced in surface (5-15 cm; P) and deeper (40-50 cm: P and Mg) soils for symptomatic trees. Total N was also lower at 40-50 cm depth for symptomatic trees, although this did not translate into differences in plant-available nitrogen (KCl extractable NO_3_^−^ and NH_4_^+^). Soil organic matter (40-50 cm depth) content was also lower for symptomatic trees. Previous studies that have examined soil nutrient properties in relation to AOD symptom status have focussed solely on total carbon and nitrogen and analysis of rhizosphere soil (sampled to 30 cm depth) and did not detect any differences in total C or N concentration between *Q. robur* trees symptomatic and non-symptomatic for AOD at three UK sites (Pinho, D. *et al*., 2020). However Scarlett *et al*. (2021) identified relationships between nitrogen cycling microbial communities and oak health status. Analysis of soils in relation to oak decline in non-AOD contexts have found reductions in total N and extractable concentrations of P, Mg Ca, K and Na, but only beneath dead *Q. robur* (Rozas & Sampedro, 2013). In Mediterranean forests, significant reductions in available P in soil have also been reported for *Q. suber* affected by *Phytophthora cinnamomi*-induced dieback (Avila *et al*., 2016). In contrast (drought-induced) crown defoliation in *Q. ilex* has been associated with accumulation of available P, N and soil organic carbon in surface (0-10 cm) soils, but, also a net loss of soil available Mg^2+^ (García-Angulo *et al*., 2020).

#### 4.2.2 Potential role as stressors in the AOD decline spiral

Water saturation presents a significant stressor for many tree species, with risk of soil hypoxia/anoxia and accumulations of gaseous metabolic products and reduced soil chemical species such as ferrous iron. These conditions can severely disrupt growth and physiological processes in tree species that lack specific adaptations to waterlogging stress, potentially resulting in dieback of affected tissues (Kreuzwieser & Rennenberg, 2014). However, *Q. robur* and, to a lesser extent, *Q. petraea* exhibit a relatively high tolerance to waterlogging and root hypoxia. These species have demonstrated an ability to adjust root growth and root-to-shoot biomass ratios to adapt to waterlogging constraints (Schmull & Thomas, 2000). While the response of conifers to Fe(II) toxicity caused by waterlogged conditions in peaty soils has been studied (Sanderson & Armstrong, 1980), the possibility of waterlogging-induced Fe(II) toxicity on oaks growing in mineral soils has not, to our knowledge, been considered previously.

Previous studies suggest that oak decline may not be attributable solely to waterlogging stress but rather to a complex interplay of factors. Hypotheses propose that the presence of a high-water table from late autumn through spring, combined with summer droughts, might exacerbate oak decline. This seasonal variation in water availability creates sharp differentials that could contribute to tree stress (Oosterbaan & Nabuurs, 1991; Thomas & Hartmann, 1996; Thomas & Hartmann, 1998; Vincke & Delvaux, 2005). During summer, decreasing soil moisture can lead to soil shrinkage and crack formation, which, in turn, promotes bypass flow and increases drought risk. This risk is compounded by potential waterlogging-induced impairments in root function from previous seasons (Thomas & Hartmann, 1998; Vincke & Delvaux, 2005). It is possible that the symptomatic trees in our study experienced greater impairments to root systems given the shallower depth to seasonal water saturation, which diminished water (and nutrient) uptake capacity in summer, accentuating summer drought as a predisposing stressor for AOD.

The ESC (Pyatt et al. 2001) estimates the three sites to be suitable for both *Quercus petraea* and *Q. robur* but the suitability estimate is based on average climatic conditions for the period 1981 to 2010 and does not account for site conditions affected by extreme dry summers and wetter winter conditions as have occurred over the last decade. Furthermore, reduced availability of essential nutrients such as Mg and P, as found for symptomatic trees in our study, may also play a role in the predisposition to oak decline. In general optimal P supply promotes disease resistance in plants (Datnoff *et al*., 2023), and, while magnesium’s role in pathogenesis is less well-documented (Huber *et al*., 2012), it is known to have many physiological and structural roles that may influence disease resistance (Huber & Jones, 2013). European-wide surveys indicate that oak nutrition, particularly for *Q. petraea*, is often deficient in P (Jonard *et al*., 2015), and Mg deficiencies are also common in European forests (Armbruster *et al*., 2002). Additionally, areas affected by elevated N deposition, which exceeds critical loads, have been linked to oak decline (Brown *et al*., 2018). Excess nitrogen can disrupt nutrient balance, potentially leading to relative deficiencies of P and Mg, which may exacerbate decline. Given the observed decline in absolute or relative P and Mg nutrition in oaks, and assuming the ‘available’ or ‘exchangeable’ concentrations in the soil reflect actual availability to roots, it is possible that P and Mg deficiencies contributed as predisposing factors for AOD. Foliar analysis would be necessary to confirm whether these deficiencies are present in the AOD-symptomatic oaks studied here. However, while nutrient imbalances in oak stands have been widely reported across Europe (Thomas *et al*., 2002), and in some cases linked to oak mortality (Morillas *et al*., 2012) or foliage health (Thomas & Büttner, 1998), many studies have found no strong association between nutrient status and oak vigour (Thomas *et al*., 2002). Building on these findings in broader oak decline research, our study offers novel evidence specifically linking soil nutrient imbalances, such as reduced Mg and P, and water balance indicators like elevated Fe and gleying, to AOD symptoms.

#### 4.2.3 Potential role as indicators of health status

While variation in soil physico-chemical properties discussed above could play a predisposing role in AOD, it is equally possible that the recorded variation in soil conditions is a consequence of the health status of the tree, reflecting tree-environment feedbacks. As discussed (section 4.1), trees with AOD symptoms may have reduced photosynthesis and leaf/root litter input to soil and impaired water and nutrient uptake, thereby influencing soil properties. This potential feedback complicates the interpretation of predisposing cause and effect in studies of AOD.

Previous research has reported higher soil moisture in the root zones of declining or diseased trees compared to non-affected trees. For instance, studies on *Quercus ilex* and holm oak have shown that declining trees exhibit higher soil moisture, likely due to reduced transpiration (Corcobado *et al*., 2013; Rodríguez *et al*., 2023). However, these effects can vary depending on the stage of decline and the climatic conditions in a given year (Rodríguez *et al*., 2023). Similarly, in stands of balsam fir and black spruce, insect defoliation has been associated with increased soil water content, as defoliated trees alter the water balance by shifting the ratio of water input (precipitation) to output (evapotranspiration) (Balducci *et al*., 2020). Conceptually, it has been hypothesized that increased soil moisture following mountain pine beetle attacks could further impact soil and ecosystem biogeochemical processes, particularly carbon and nitrogen cycling (Edburg *et al*., 2012).

Previous meta-analyses have established that biotic disturbances, including pest and pathogen impacts, significantly reduce soil organic carbon (SOC) concentrations (Zhang *et al*., 2015). This reduction is primarily due to decreased photosynthetic carbon inputs from both above-ground litter and below-ground root activity (Holden & Treseder, 2013). In our study, reduction in soil organic matter (and associated SOC) content for AOD symptomatic trees was only significant for samples taken from 40-50 cm. In the absence of significant earthworm activity and thus incorporation of surface litter to depth, a difference in SOM at depth is likely a signature of reduced C inputs via roots and rhizodeposition by symptomatic trees. Strong winds may partially homogenize the distribution of surface litter across the forest floor accounting for the lack of an effect at 5-15 cm, whereas at 40-50 cm depth, the reduction in inputs in root-derived carbon would be not subject to the same spatial homogenization. In addition, when compared to above-ground litter, root-derived carbon is thought to make greater contributions to SOM accumulation due to greater potential for inputs to be protected from decomposition through stabilizing interactions with soil minerals at depth (Jackson *et al*., 2017). Asymptomatic trees, in our study, were associated with higher silt at depth which might also have contributed to higher OM through mineral protection. Thus, due to protection from both homogenization by wind and decomposition via mineral associations, SOM content at depth may be more sensitive to reductions in photosynthetic C input for symptomatic trees.

Among the studies that have examined associations between soil nitrogen and AOD (Pinho, Diogo *et al*., 2020) or oak declines (Rozas & Sampedro, 2013), the research by Thomas and Buttner (1998) stands out for consideration of the consequences of oak health on soil nitrogen budgets. Their study in northwestern Germany suggested that in declining oak stands, reduced root uptake may lead to nitrogen losses through increased leaching, indicating that nitrogen loss is a consequence, rather than a cause, of oak decline. Our findings of reduced total nitrogen at depth align with the hypothesis that nitrogen is being lost from soils surrounding AOD-affected oaks. With reduced nitrogen uptake, more nitrogen remains in forms susceptible to loss through processes such as leaching and denitrification. Interestingly, while previous research has not directly linked denitrification gene abundance to tree health (Scarlett *et al*., 2021), possible greater seasonal water saturation around our symptomatic trees —evidenced by gleying and elevated iron concentrations— could create reducing conditions favourable for denitrification.

Thomas and Buttner (1998) also noted that reduced nitrogen uptake and increased nitrate output were associated with significant magnesium loss. In our study, symptomatic trees were similarly linked to lower soil Mg levels, with the greatest reduction in N observed in areas with the most significant Mg depletion. This supports the idea that Mg leaching occurs preferentially, as cations are leached alongside anions like nitrate to maintain electrical neutrality in the soil solution (Barber, 1995).

Phosphorus cycling in forest ecosystems is typically characterized as a “closed cycle,” where phosphorus pools within soil and biomass are substantially larger than the fluxes through atmospheric deposition, weathering, leaching, or harvesting (Ilg *et al*., 2009). Given this closed cycle, possible reductions in phosphorus input through reduced litterfall, due to tree decline, might be counterbalanced by reductions in phosphorus uptake, maintaining a degree of equilibrium. However, in our study, we observed that reductions in soil available phosphorus, particularly in the 5-15 cm depth, were linked to symptomatic trees. This depth is likely influenced by above-ground litter inputs, and reductions in litterfall from declining trees might, in our case, have influenced available phosphorus levels.

Leaf litter manipulation experiments have shown varied effects on soil phosphorus concentrations. When compared to non-manipulated (control) litter inputs, litter addition has been shown to increase soil available P in the A horizon (Huang & Spohn, 2015). This finding is consistent with increased P input as litter and subsequent microbial P minerlization. In contrast, litter exclusion also increased available P in the A horizon relative to the control (Huang & Spohn, 2015) which was attributed to a rise in fine root growth in this horizon in the absence of surface litter and rhizosphere-enhanced P mineralization acitivty. This indicates that while litter contributes to P cycling, roots play a crucial role in regulating phosphorus redistribution and availability in soil. Avila *et al*. (2016) reported significant reductions in available phosphorus in soils affected by *Phytophthora cinnamomi*-induced *Q. suber* dieback. They attributed these reductions to decreased root activity in phosphorus mineralization and solubilization, due to reduced phosphatase production and organic acid exudation. Our findings align with this observation, suggesting that symptomatic trees with AOD may exhibit reduced root activity, impairing their ability to mobilize and solubilize phosphorus from unavailable sources.

Additionally, our supplementary analyses (Supplementary Fig. S3) indicated that not only available phosphorus but also total phosphorus concentrations were reduced in soils associated with AOD-symptomatic trees. This could suggest that symptomatic trees are affecting both the proportion of available phosphorus and the total phosphorus pool in the soil. Trees experiencing health declines might tighten internal nutrient cycling, redistributing nutrients from senescing leaves to stem tissues during dormancy. This internal cycling is a known strategy in temperate deciduous species to conserve nutrients when faced with deficiencies (Achat *et al*., 2018), regulated by soil nutrient availability (Netzer *et al*., 2017; Achat *et al*., 2018). Thus, symptomatic trees may exhibit reduced litterfall quality, which could further affect soil phosphorus availability.

Finally, the evidence (gleying, elevated available iron) that AOD symptomatic trees may experience more intense seasonal waterlogging might also be relevant to interpretations since phosphorus availability is influenced by interactions with iron (Burgin *et al*., 2011). During waterlogging, Fe is reduced from Fe(III) to Fe(II), temporarily releasing phosphorus bound to secondary minerals, such as iron (hydr-)oxides. A greater solubility of P under reducing conditions could lead to loss of P via leaching. However, when the soil dries and oxygen returns, Fe(II) reoxidizes to Fe(III), re-forming reactive iron (hydr-)oxides that immobilize phosphorus, reducing its extraction by the Olsen P method used. Fluctuating redox conditions for AOD symptomatic trees could thus contribute to depleted total and available P pools via effects on P-Fe interactions (Miller *et al*., 2001) which might impact tree P uptake (Thomas & Buttner, 1998).

### 4.3 Synthesis of cause and effect

The preceding discussion argues how water excess and wet-dry extremes, elevated iron, and reduced concentrations of other essential nutrients could potentially stress trees (section 4.2.2). While water and nutrient factors have been linked to oak decline in past studies, their specific role in AOD at a local scale has not been clearly established. The predisposing, inciting, and contributing factors of forest decline are conceptualized as operating in a specific sequence (Fig. 6a), with predisposing factors relating to soil and inherent environmental factors acting early in the decline process (Manion & Lachance, 1992). It is conceivable that the observed variations in soil properties at the tree scale reflect inherent local heterogeneity, which interacts with inciting factors to influence tree susceptibility to pathogens.

**Figure 6.**
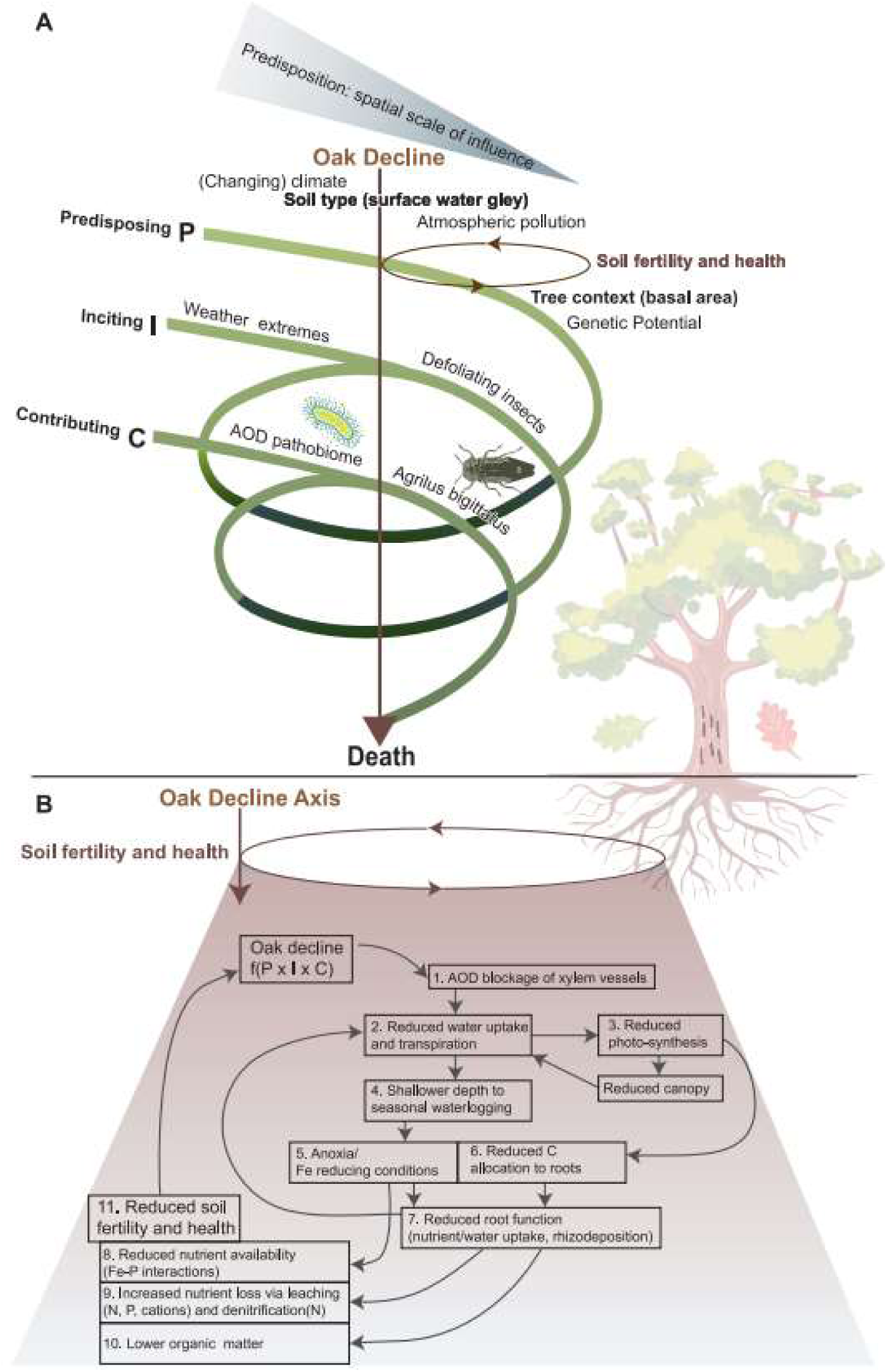
(A) Conceptual model illustrating the decline disease spiral model (adapted from Denman *et al*. (2022) with permission from Elsevier) for acute oak decline (AOD), highlighting the interactions between predisposing, inciting, and contributing factors. Predisposing factors represent chronic, long-term influences that weaken a tree’s resilience and create a vulnerable baseline. The wedge illustrates how these factors operate across both broad (e.g., climate) and more localized (e.g., individual tree characteristics) spatial scales. This study identifies soil type interacting with individual tree context (highlighted in bold) as a potentially significant predisposing factors. Inciting factors are short-term stressors, such as extreme weather events (e.g., drought) or insect defoliation, which may trigger visible symptoms of decline in trees already weakened by predisposing factors. Contributing factors are secondary agents or conditions, such as pathogens (here, AOD-associated bacteria) and pests (here, possibly Agrilus biguttatus), that exploit weakened trees and accelerate decline, potentially leading to mortality. The negative feedback hypothesized in this study between the oak health axis (arrow) and soil fertility/health is depicted in brown, further detailed in (B). (B) Expanded view of the hypothesized feedback mechanisms between oak health and soil fertility and health in the context of AOD. Oak decline is conceptualized as a function of interactions between predisposing (P), inciting (I), and contributing factors (C). For AOD, bacterial infection and xylem disruption [1] impair water transport [2], reducing stomatal conductance and photosynthesis [3]—similar to drought stress. Reduced water uptake leads to locally high soil moisture during periods of elevated water tables (e.g., in spring), leading to a shallower depth to seasonal waterlogging [4] from a perched water table (surface water gley), exposing a larger portion of the root system to anoxic, iron-reducing conditions [5], which can reduce phosphorus (P) availability through Fe-P redox reactions [8]. Reduced carbon allocation to root systems [6] further diminishes root function, limiting water and nutrient uptake [7] and increasing nutrient losses through leaching and gaseous emissions [9]. Lower carbon inputs to soil via reduced rhizodeposition decrease soil organic matter [10] and soil health. These overall changes in soil fertility and health [11] create a reinforcing negative feedback loop that exacerbates oak health decline.

Fundamentally, spatial variations in soil properties arise from the interplay of parent material, climate, topography, and biotic factors, assuming a constant time for soil formation (Jenny, 1941). Within a woodland site, individual trees experience similar broad climatic conditions, but local differences in soil properties might reflect variations in soil parent material, micro-topography or biological context. There was no evidence to suggest that differences in microtopography played a significant role in local variations at the tree scale, as indicated by the lack of difference in Compound Topographic Index. Although soils for AOD-affected trees had lower silt content (indicating potential variation in surficial parent materials), these variations were small compared to the larger textural differences observed between sites. However, in our sample, symptomatic trees were located in areas with a lower basal area within a 20 m radius. This means that these trees were surrounded by fewer or smaller neighbouring trees, compared to the asymptomatic ones. While the reasons for this difference in context remain unknown, it implies that AOD-affected trees may currently experience less competition from other trees, although increased competition from ground vegetation is possible (Henneron *et al*., 2017).

Despite this, the lower basal area could explain the water balance indicators identified earlier, as clusters of trees with reduced stem density might collectively be less effective at removing water from the root zone via transpiration (del Campo *et al*., 2022). Such altered water dynamics, especially when water tables are high on a seasonal basis, could exacerbate susceptibility to water-related stresses. The context of trees with respect to lower basal area with effects on shallower soil depth to seasonal waterlogging could therefore represent a locally-acting predisposition factor for AOD. This conclusion is put forward as it is difficult to envisage a scheme whereby the reverse explanation - that the lower basal area is *a result* of AOD – could be plausible. The possibility that our method of tree selection inadvertently focused on AOD-affected trees with lower basal area cannot be entirely ruled out, but it seems unlikely. Further testing at other sites would be necessary to determine if lower basal area around symptomatic trees consistently correlates with AOD.

In addition to exploring cause-and-effect relationships for tree context factors, such as basal area, the evidence regarding whether the recorded soil variations are predisposing factors or consequences of tree health needs further evaluation. The temporal aspect of tree decline makes it challenging, as measurements taken at a single time point attempt to reconstruct past events and understanding soil variability prior to tree establishment is not possible (Aponte *et al*., 2013). The spatial distribution of trees has been shown to drive spatial heterogeneity of soil resources by varying litter fall inputs (Andivia *et al*., 2015). As discussed (section 4.2.3), soil chemical signatures beneath trees with AOD symptoms may be explainable by tree responses to the disease such as reduced photosynthesis, impaired nutrient and water uptake, creating a feedback to their own soil environment. If this feedback exists, our evidence suggests that it is a negative loop (Fig. 6b), where the decline in tree health worsens the surrounding soil health through reduced SOM, nutrient loss, waterlogging, and increased Fe levels, potentially leading to toxicity. These soil conditions then exacerbate tree decline, setting up a self-reinforcing cycle of deterioration.

## Conclusions

Our study highlights that trees exhibiting AOD symptoms are linked to tree-scale soil water and nutrient properties, but we lack definitive evidence to determine whether these factors are predominantly predisposing conditions or consequences of declining health. This uncertainty complicates our understanding of AOD. In this context, the decline disease spiral model could be expanded to incorporate potential feedback loops between tree health and soil conditions (Fig 6). Furthermore, the differentiation between large-scale environmental factors and localized tree-specific conditions is critical. While factors such as the likeliness of seasonal waterlogging may be linked to broader regional soil types or climatic conditions, our findings suggest that local context— such as the local scale (0-20 m) basal area of surrounding trees or the micro-scale soil properties— could play a crucial role in individual tree responses. Although tree genetic potential was not investigated in this study, it remains an additional factor that could contribute to individual tree responses to AOD.

Future research should focus on disentangling cause-effect relationships through long-term monitoring, experiments and mechanistic modelling to better understand how soil conditions and tree health interact over time, and whether internal feedback mechanisms significantly contribute to the AOD decline spiral. A key gap is an understanding of how tree health changes to enable the bacterial pathogens to proliferate and cause bleeding cankers. Ultimately, woodland management in the context of AOD must be informed by a clearer mechanistic understanding. However, adaptive strategies such as promoting natural regeneration to favour phenotypes more able to acclimate to changes in the seasonal climate (Cavers & Cottrell, 2014; Ennos, 2014), addressing local soil nutrient imbalances and seasonal water extremes may help build resilience in oak woodlands and reduce AOD impacts and feedbacks.

## Acknowledgements

The work was funded by Defra through the BBSRC grant Protecting Oak Ecosystems (PuRpOsE): BB/N022831/1. We thank Anne Dudley, Ilse Kamerling, Chris Speed, Jessica Princivalle, Harriet Robson, Shyamali Roy, Anna Oliver, Andrew Rattey, Stuart A’Hara, Liz Richardson and the Forest Research Field Data Services team for help with sampling and sample analyses. We thank Kelvin Balcombe for help with statistical analysis, Joan Cottrell for help with oak tree species identification and Sarah Lambert-Gates for help with illustrations in Fig. 6. We thank the Woodland Trust for access to Stratfield Brake and Monks Wood as well as additional funding to RWJ, DR and AB to support the analysis of oak trees in Stratfield Brake. We thank Writtle Forest Limited for access to sites.

## Data availability

The data generated in this study are in the process of being deposited in the UK Natural Environment Research Council’s Environmental Information Data Centre (EIDC) (https://eidc.ac.uk/) with the following data identifier: 9947ccc9-7396-4821-b095-c374bd857305.

## References

Achat DL, Pousse N, Nicolas M, Augusto L. 2018. Nutrient remobilization in tree foliage as affected by soil nutrients and leaf life span. Ecological Monographs 88(3): 408–428.

Andivia E, Fernández M, Alejano R, Vázquez-Piqué J. 2015. Tree patch distribution drives spatial heterogeneity of soil traits in cork oak woodlands. Annals of Forest Science 72(5): 549–559.

Aponte C, García LV, Marañón T. 2013. Tree species effects on nutrient cycling and soil biota: A feedback mechanism favouring species coexistence. Forest Ecology and Management 309: 36–46.

Armbruster M, MacDonald J, Dise NB, Matzner E. 2002. Throughfall and output fluxes of Mg in European forest ecosystems: a regional assessment. Forest Ecology and Management 164(1): 137–147.

Avery B. 1980. Soil Classification for England and Wales (Higher Categories). Soil Survey Technical Monograph No.14. Harpenden: Rothamsted Experimental Station.

Avery B. 1990. Soils of the British Isles. Wallingford: CABI.

Avila JM, Gallardo A, Ibáñez B, Gómez-Aparicio L. 2016. Quercus suber dieback alters soil respiration and nutrient availability in Mediterranean forests. Journal of Ecology 104(5): 1441–1452.

Baldrian P. 2014. Distribution of Extracellular Enzymes in Soils: Spatial Heterogeneity and Determining Factors at Various Scales. Soil Science Society of America Journal 78(1): 11–18.

Balducci L, Fierravanti A, Rossi S, Delzon S, De Grandpré L, Kneeshaw DD, Deslauriers A. 2020. The paradox of defoliation: Declining tree water status with increasing soil water content. Agricultural and Forest Meteorology 290.

Barber S. 1995. Soil Nutrient Bioavailability: A Mechanistic Approach: John Wiley & Sons.

Barsoum N, A’Hara SW, Cottrell JE, Forster J, Garcia MSJ, Schonrogge K, Shaw L. 2021. Root ectomycorrhizal status of oak trees symptomatic and asymptomatic for Acute Oak Decline in southern Britain. Forest Ecology and Management 482: 118800.

Beatty SW. 1984. Influence of microtopography and canopy species on spatial patterns of forest understory plants. Ecology 65(5): 1406–1419.

Booth O. 2021. Consideration of pathogenicity of bacterial species associated with bleed canker on Quercus sp. and evaluation of potential methods of control. PhD Thesis, University of Reading Reading, U.K.

Brady C, Denman S, Kirk S, Venter S, Rodriguez-Palenzuela P, Coutinho T. 2010. Description of *Gibbsiella quercinecans* gen. nov., sp. nov., associated with Acute Oak Decline. Systematic and Applied Microbiology 33(8): 444–450.

Brady C, Hunter G, Kirk S, Arnold D, Denman S. 2014. *Rahnella victoriana* sp nov., *Rahnella bruchi* sp nov., *Rahnella woolbedingensis* sp nov., classification of *Rahnella* genomospecies 2 and 3 as *Rahnella variigena* sp nov and *Rahnella inusitata* sp nov., respectively and emended description of the genus *Rahnella*. Systematic and Applied Microbiology 37(8): 545–552.

Brady CL, Cleenwerck I, Denman S, Venter SN, Rodriguez-Palenzuela P, Coutinho TA, De Vos P. 2012. Proposal to reclassify *Brenneria quercina* (Hildebrand and Schroth 1967) Hauben et al. 1999 into a new genus, *Lonsdalea* gen. nov., as *Lonsdalea quercina* comb. nov., descriptions of *Lonsdalea quercina* subsp *quercina* comb. nov., *Lonsdalea quercina* subsp *iberica* subsp nov and *Lonsdalea quercina* subsp *britannica* subsp nov., emendation of the description of the genus *Brenneria*, reclassification of *Dickeya* dieffenbachiae as Dickeya *dadantii* subsp *dieffenbachiae* comb. nov., and emendation of the description of *Dickeya dadantii*. International Journal of Systematic and Evolutionary Microbiology 62: 1592–1602.

Brady N, Weil R. 2013. The Nature and Properties of Soils. Harlow, U.K.: Pearson Education Limited.

Broberg M, Doonan J, Mundt F, Denman S, McDonald JE. 2018. Integrated multi-omic analysis of host-microbiota interactions in acute oak decline. Microbiome 6(1): 21.

Brown N, Jeger M, Kirk S, Xu X, Denman S. 2016. Spatial and temporal patterns in symptom expression within eight woodlands affected by Acute Oak Decline. Forest Ecology and Management 360: 97–109.

Brown N, van den Bosch F, Parnell S, Denman S. 2017. Integrating regulatory surveys and citizen science to map outbreaks of forest diseases: acute oak decline in England and Wales. Proceedings of the Royal Society B-Biological Sciences 284(1859).

Brown N, Vanguelova E, Parnell S, Broadmeadow S, Denman S. 2018. Predisposition of forests to biotic disturbance: Predicting the distribution of Acute Oak Decline using environmental factors. Forest Ecology and Management 407: 145–154.

Burgin AJ, Yang WH, Hamilton SK, Silver WL. 2011. Beyond carbon and nitrogen: how the microbial energy economy couples elemental cycles in diverse ecosystems. Frontiers in Ecology and the Environment 9(1): 44–52.

Cambon MC, Thomas G, Caulfield J, Crampton M, Reed K, Doonan JM, Hussain U, Denman S, Vuts J, McDonald JE. 2024. Chemical cues from beetle larvae trigger proliferation and putative virulence gene expression of a plant pathogen. bioRxiv: 2023.2011.2021.568124.

Cavers S, Cottrell JE. 2014. The basis of resilience in forest tree species and its use in adaptive forest management in Britain. Forestry: An International Journal of Forest Research 88(1): 13–26.

Cochard H, Breda N, Granier A. 1996. Whole tree hydraulic conductance and water loss regulation in Quercus during drought: evidence for stomatal control of embolism? Annales Des Sciences Forestieres 53: 197–206.

Cools N, De Vos B 2016. Part X: Sampling and Analysis of Soil. In: UNNECE ICP Forrests Programme Co-ordinating Centre (Eds). Manual on methods and criteria for harmonized sampling, assessment, monitoring and analysis of the effetcs of air pollution on forests. Eberswalde, Germany: Thünen Institute of Forest Ecosystems.

Corcobado T, Cubera E, Moreno G, Solla A. 2013. *Quercus ilex* forests are influenced by annual variations in water table, soil water deficit and fine root loss caused by *Phytophthora cinnamomi*. Agricultural and Forest Meteorology 169: 92–99.

Crampton B, Brady C, Denman S 2022. Bacterial Tree Disease Fact Sheets—Acute oak decline.

Datnoff LE, Elmer WH, Rodrigues FA, Huber DM. 2023. Mineral nutrition and plant disease. St. Paul, Minnesota, U.S.A: The American Phytopathological Society.

del Campo AD, Otsuki K, Serengil Y, Blanco JA, Yousefpour R, Wei X. 2022. A global synthesis on the effects of thinning on hydrological processes: Implications for forest management. Forest Ecology and Management 519: 120324.

Denman S, Brady C, Kirk S, Cleenwerck I, Venter S, Coutinho T, De Vos P. 2012. *Brenneria goodwinii* sp nov., associated with acute oak decline in the UK. International Journal of Systematic and Evolutionary Microbiology 62: 2451–2456.

Denman S, Brown N, Kirk S, Jeger M, Webber J. 2014. A description of the symptoms of Acute Oak Decline in Britain and a comparative review on causes of similar disorders on oak in Europe. Forestry 87(4): 535–551.

Denman S, Brown N, Vanguelova E, Crampton B 2022. Chapter 14 - Temperate Oak Declines: Biotic and abiotic predisposition drivers. In: Asiegbu FO, Kovalchuk A eds. *Forest Microbiology*: Academic Press, 239–263.

Denman S, Doonan J, Ransom-Jones E, Broberg M, Plummer S, Kirk S, Scarlett K, Griffiths AR, Kaczmarek M, Forster J, et al. 2018. Microbiome and infectivity studies reveal complex polyspecies tree disease in Acute Oak Decline. Isme Journal 12(2): 386–399.

Denman S, Plummer S, Kirk S, Peace A, McDonald JE. 2016. Isolation studies reveal a shift in the cultivable microbiome of oak affected with Acute Oak Decline. Systematic and Applied Microbiology 39(7): 484–490.

Denman S, Webber J. 2009. Oak declines: new definitions and new episodes in Britain. Quarterly Journal of Forestry 103(4): 285–290.

Doonan J, Denman S, Pachebat JA, McDonald JE. 2019. Genomic analysis of bacteria in the Acute Oak Decline pathobiome. Microbial Genomics 5(1).

Edburg SL, Hicke JA, Brooks PD, Pendall EG, Ewers BE, Norton U, Gochis D, Gutmann ED, Meddens AJ. 2012. Cascading impacts of bark beetle-caused tree mortality on coupled biogeophysical and biogeochemical processes. Frontiers in Ecology and the Environment 10(8): 416–424.

Ennos RA. 2014. Resilience of forests to pathogens: an evolutionary ecology perspective. Forestry: An International Journal of Forest Research 88(1): 41–52.

Farley RA, Fitter AH. 1999. Temporal and spatial variation in soil resources in a deciduous woodland. Journal of Ecology 87(4): 688–696.

Fernandes C, Duarte L, Naves P, Sousa E, Cruz L. 2022. First report of *Brenneria goodwinii* causing acute oak decline on *Quercus suber* in Portugal. Journal of Plant Pathology 104(2): 837–838.

Flinn KM, Marks PL. 2007. Agricultural legacies in forest environments: Tree communities, soil properties, and light availability. Ecological Applications 17(2): 452–463.

García-Angulo D, Hereş AM, Fernández-López M, Flores O, Sanz MJ, Rey A, Valladares F, Curiel Yuste J. 2020. Holm oak decline and mortality exacerbates drought effects on soil biogeochemical cycling and soil microbial communities across a climatic gradient. Soil Biology and Biochemistry 149: 107921.

Gonzalez AJ, Ciordia M. 2020. *Brenneria goodwinii* and *Gibbsiella quercinecans* isolated from weeping cankers on *Quercus robur L.* in Spain. European Journal of Plant Pathology 156(3): 965–969.

González AJ, Ciordia M. 2020. Brenneria goodwinii and Gibbsiella quercinecans isolated from weeping cankers on Quercus robur L. in Spain. European Journal of Plant Pathology 156(3): 965–969.

Gosling RH, Jackson RW, Elliot M, Nichols CP. 2024. Oak declines: Reviewing the evidence for causes, management implications and research gaps. Ecological Solutions and Evidence 5(4): e12395.

Gotoh S, Patrick Jr. WH. 1974. Transformation of Iron in a Waterlogged Soil as Influenced by Redox Potential and pH. Soil Science Society of America Journal 38(1): 66–71.

Greenwell B, Boehmke B, Cunningham J, Developers G 2020. Generalized Boosted Regressin Models. R package version 2.1.8. https://CRAN.R-project.org/package=gbm.

Guichoux E, Garnier-Géré P, Lagache L, Lang T, Boury C, Petit RJ. 2013. Outlier loci highlight the direction of introgression in oaks. Mol Ecol 22(2): 450–462.

Hamilton G 1975. Forest Mensuration Handbook (Forestry Commission Booklet no. 39). London: Stationary Office Books.

Hardoim PR, van Overbeek LS, Berg G, Pirttila AM, Compant S, Campisano A, Doering M, Sessitsch A. 2015. The Hidden World within Plants: Ecological and Evolutionary Considerations for Defining Functioning of Microbial Endophytes. Microbiology and Molecular Biology Reviews 79(3): 293–320.

Hartig F 2020. DHARMa: Residual Diagnostics for Hierarchical (Multi-Level / Mixed) Regression Models. R package version 0.3.3.0. https://CRAN.R-project.org/package=DHARMa.

Henneron L, Chauvat M, Archaux F, Akpa-Vinceslas M, Bureau F, Dumas Y, Mignot L, Ningre F, Perret S, Richter C, et al. 2017. Plant interactions as biotic drivers of plasticity in leaf litter traits and decomposability of Quercus petraea. Ecological Monographs 87(2): 321–340.

Holden SR, Treseder KK. 2013. A meta-analysis of soil microbial biomass responses to forest disturbances. Frontiers in Microbiology 4.

Houston D 1992. A host-stress-saprogen model for forest dieback -decline diseases. In: Manion P, Lachance D eds. Forest Decline Concepts. St. Paul, Minnesota, USA.: APS Press, 3–25.

Huang W, Spohn M. 2015. Effects of long-term litter manipulation on soil carbon, nitrogen, and phosphorus in a temperate deciduous forest. Soil Biology & Biochemistry 83: 12–18.

Huber D, Jones J. 2013. The role of magnesium in plant disease. Plant and Soil 368: 73–85.

Huber D, Römheld V, Weinmann M 2012. Chapter 10 - Relationship between Nutrition, Plant Diseases and Pests. In: Marschner P ed. *Marschner’s Mineral Nutrition of Higher Plants (Third Edition)*: Academic Press, 283–298.

Ilg K, Wellbrock N, Lux W. 2009. Phosphorus supply and cycling at long-term forest monitoring sites in Germany. European Journal of Forest Research 128(5): 483–492.

Jackson RB, Lajtha K, Crow SE, Hugelius G, Kramer MG, Piñeiro G 2017. The Ecology of Soil Carbon: Pools, Vulnerabilities, and Biotic and Abiotic Controls. In: Futuyma DJ ed. Annual Review of Ecology, Evolution, and Systematics, Vol 48, 419–445.

Jenny H. 1941. Factors of Soil Formation: A System of Quantitative Pedology. New York: Dover Publications.

Jonard M, Fuerst A, Verstraeten A, Thimonier A, Timmermann V, Potocic N, Waldner P, Benham S, Hansen K, Merila P, et al. 2015. Tree mineral nutrition is deteriorating in Europe. Global Change Biology 21(1): 418–430.

Jonsson U, Jung T, Sonesson K, Rosengren U. 2005. Relationships between health of Quercus robur, occurrence of Phytophthora species and site conditions in southern Sweden. Plant Pathology 54(4): 502–511.

Kandel SL, Firrincieli A, Joubert PM, Okubara PA, Leston ND, McGeorge KM, Mugnozza GS, Harfouche A, Kim SH, Doty SL. 2017. An In vitro Study of Bio-Control and Plant Growth Promotion Potential of Salicaceae Endophytes. Frontiers in Microbiology 8: 16.

Kint V, Vansteenkiste D, Aertsen W, De Vos B, Bequet R, Van Acker J, Muys B. 2012. Forest structure and soil fertility determine internal stem morphology of Pedunculate oak: a modelling approach using boosted regression trees. European Journal of Forest Research 131(3): 609–622.

Kremer A, Dupouey J-L, Deans J, Cottrell J, Csaikl U, Finkeldey R, Espinel S, Jensen J, Kleinschmit J, Dam B, et al. 2002. Leaf morphological differentiation between *Quercus robur* and *Quercus petraea* is stable across western European mixed oak stands. Annals of Forest Science 59: 777–787.

Kreuzwieser J, Rennenberg H. 2014. Molecular and physiological responses of trees to waterlogging stress. Plant Cell and Environment 37(10): 2245–2259.

Kuhn M 2021. caret: Classification and Regression Training. R package version 6.0-90. https://CRAN.R-project.org/package=caret.

Lakatos F, Mirtchev S, Mehmeti A, Shabanaj H 2014. Manual for visual assessment of forest crown condition. Pristina: UN FAO.

Lenth R 2021. emmeans: Estimated Marginal Means, aka Least-Squares Means. R package version 1.6.2–1. https://CRAN.R-project.org/package=emmeans.

Maddock D, Brady C, Denman S, Arnold D. 2023. Bacteria Associated with Acute Oak Decline: Where Did They Come From? We Know Where They Go. Microorganisms 11(11): 2789.

Maes M, Baeyen S, De Croo H, De Smet K, Steenackers M. 2002. Monitoring of endophytic *Brenneria salicis* in willow and its relation to watermark disease. Plant Protection Science 38(Special Issue 2): 528–530.

Maes M, Huvenne H, Messens E. 2009. Brenneria salicis, the bacterium causing watermark disease in willow, resides as an endophyte in wood. Environmental Microbiology 11(6): 1453–1462.

Manion P, Lachance D 1992. Forest decline concepts: An overview. In: Manion P, Lachance D eds. Forest Decline Concepts. St. Paul, Minnesota, USA.: APS Press, 181–190.

Miles J 1986. What are the effects of trees on soils? In: Jenkins D. e ed. Trees and wildlife in the Scottish uplands. NERC/ITE, 55 - 62. (ITE Symposium, 17), NERC, URL: http://nora.nerc.ac.uk/5293/

Miller AJ, Schuur EAG, Chadwick OA. 2001. Redox control of phosphorus pools in Hawaiian montane forest soils. Geoderma 102(3-4): 219–237.

Moradi-Amirabad Y, Rahimian H, Babaeizad V, Denman S. 2019. *Brenneria spp*. and *Rahnella victoriana* associated with acute oak decline symptoms on oak and hornbeam in Iran. Forest Pathology 49(4).

Morillas L, Gallardo A, Portillo-Estrada M, Covelo F. 2012. Nutritional status of Quercus suber populations under contrasting tree dieback. Forestry 85(3): 369–377.

Netzer F, Schmid C, Herschbach C, Rennenberg H. 2017. Phosphorus-nutrition of European beech (*Fagus sylvatica* L.) during annual growth depends on tree age and P-availability in the soil. Environmental and Experimental Botany 137: 194–207.

Olsen S, Cole C, Watanabe F, Dean L 1954. Estimation of available phosphorus in soils by extraction with sodium bicarbonate. USDA Circular 939. Washington DC: USDA.

Oosterbaan A, Nabuurs GJ. 1991. Relationship between oak decline and groundwater class in the Netherlands. Plant and Soil 136(1): 87–93.

Pinho D, Barroso C, Froufe H, Brown N, Vanguelova E, Egas C, Denman S. 2020. Linking Tree Health, Rhizosphere Physicochemical Properties, and Microbiome in Acute Oak Decline. Forests 11(11): 1153.

Pinho D, Barroso C, Froufe H, Brown N, Vanguelova E, Egas C, Denman S. 2020. Linking Tree Health, Rhizosphere Physicochemical Properties, and Microbiome in Acute Oak Decline. Forests 11(11).

Pyatt G, Ray D, Fletcher J 2001. An Ecological Site Classification for Forestry in Great Britain. Bulletin 124. Edinburgh: Forestry Commission.

Reed K, Denman S, Leather SR, Forster J, Inward DJG. 2018. The lifecycle of Agrilus biguttatus: the role of temperature in its development and distribution, and implications for Acute Oak Decline. Agricultural and Forest Entomology 20(3): 334–346.

Rodríguez A, Durán J, Curiel Yuste J, Valladares F, Rey A. 2023. The effect of tree decline over soil water content largely controls soil respiration dynamics in a Mediterranean woodland. Agricultural and Forest Meteorology 333: 109398.

Rosenblueth M, Martinez-Romero E. 2006. Bacterial endophytes and their interactions with hosts. Molecular Plant-Microbe Interactions 19(8): 827–837.

Rosengren U, Göransson H, Jönsson U, Stjernquist I, Thelin G, Wallander H. 2006. Functional Biodiversity Aspects on the Nutrient Sustainability in Forests-Importance of Root Distribution. Journal of Sustainable Forestry 21(2-3): 77–100.

Rozas V, Sampedro L. 2013. Soil chemical properties and dieback of Quercus robur in Atlantic wet forests after a weather extreme. Plant and Soil 373(1-2): 673–685.

Ruffner B, Schneider S, Meyer J, Queloz V, Rigling D. 2020. First report of acute oak decline disease of native and non-native oaks in Switzerland. New Disease Reports 41: 18.

Sanderson PL, Armstrong W. 1980. The responses of conifers to some of the adverse factors associated with waterlogged soils. New Phytologist 85(3): 351–362.

Sapp M, Lewis E, Moss S, Barrett B, Kirk S, Elphinstone JG, Denman S. 2016. Metabarcoding of Bacteria Associated with the Acute Oak Decline Syndrome in England. Forests 7(5): 95.

Scarlett K, Denman S, Clark DR, Forster J, Vanguelova E, Brown N, Whitby C. 2021. Relationships between nitrogen cycling microbial community abundance and composition reveal the indirect effect of soil pH on oak decline. Isme Journal 15(3): 623–635.

Schmull M, Thomas FM. 2000. Morphological and physiological reactions of young deciduous trees (Quercus robur L., Q. petraea [Matt.] Liebl., Fagus sylvatica L.) to waterlogging. Plant and Soil 225(1): 227–242.

Shen SY, Fulthorpe R. 2015. Seasonal variation of bacterial endophytes in urban trees. Frontiers in Microbiology 6: 13.

Smithwick EAH, Eissenstat DM, Lovett GM, Bowden RD, Rustad LE, Driscoll CT. 2013. Root stress and nitrogen deposition: consequences and research priorities. New Phytologist 197(3): 712–719.

Steele RC, Welch RC. 1973. Monks Wood: a nature reserve record. Huntingdon, U.K.: Nature Conservancy/Natural Environment Research Council.

Stursova M, Barta J, Santruckova H, Baldrian P. 2016. Small-scale spatial heterogeneity of ecosystem properties, microbial community composition and microbial activities in a temperate mountain forest soil. Fems Microbiology Ecology 92(12).

Thomas F. 2008. Recent advances in cause-effect research on oak decline in Europe. CAB Reviews: Perspectives in Agriculture, Veterinary Science, Nutrition and Natural Resources 3(037): 1–12.

Thomas FM, Blank R, Hartmann G. 2002. Abiotic and biotic factors and their interactions as causes of oak decline in Central Europe. Forest Pathology 32(4-5): 277–307.

Thomas FM, Buttner G. 1998. Nutrient relations in healthy and damaged stands of mature oaks on clayey soils: two case studies in northwestern Germany. Forest Ecology and Management 108(3): 301–319.

Thomas FM, Büttner G. 1998. Zusammenhänge zwischen Ernährungsstatus und Belaubungsgrad in Alteichenbeständen Nordwestdeutschlands. Forstwissenschaftliches Centralblatt vereinigt mit Tharandter forstliches Jahrbuch 117(1): 115–128.

Thomas FM, Hartmann G. 1996. Soil and tree water relations in mature oak stands of northern Germany differing in the degree of decline. Annales Des Sciences Forestieres 53(2-3): 697–720.

Thomas FM, Hartmann G. 1998. Tree rooting patterns and soil water relations of healthy and damaged stands of mature oak (Quercus robur L and Quercus petraea Matt Liebl). Plant and Soil 203(1): 145–158.

Thomas FM, Sprenger S. 2008. Responses of two closely related oak species, Quercus robur and Q. petraea, to excess manganese concentrations in the rooting medium. Tree Physiology 28(3): 343–353.

Tkaczyk M, Sikora K. 2024. The Role of Bacteria in Acute Oak Decline in South-West Poland. Microorganisms 12(5): 993.

Torres AR, Araujo WL, Cursino L, Hungria M, Plotegher F, Mostasso FL, Azevedo JL. 2008. Diversity of endophytic enterobacteria associated with different host plants. Journal of Microbiology 46(4): 373–379.

Trouve R, Bontemps JD, Collet C, Seynave I, Lebourgeois F. 2017. Radial growth resilience of sessile oak after drought is affected by site water status, stand density, and social status. Trees-Structure and Function 31(2): 517–529.

USEPA 2000. Guidance for Data Quality Assessment. U.S. Environmental Protection Agency, Washington DC.

van der Linde S, Suz LM, Orme CDL, Cox F, Andreae H, Asi E, Atkinson B, Benham S, Carroll C, Cools N, et al. 2018. Environment and host as large-scale controls of ectomycorrhizal fungi. Nature 558(7709): 243–248.

Vanguelova EI, Hirano Y, Eldhuset TD, Sas-Paszt L, Bakker MR, Puttsepp U, Brunner I, Lohmus K, Godbold D. 2007. Tree fine root Ca/Al molar ratio - Indicator of Al and acidity stress. Plant Biosystems 141(3): 460–480.

Vincke C, Delvaux B. 2005. Porosity and available water of temporarily waterlogged soils in a Quercus robur (L.) declining stand. Plant and Soil 271(1-2): 189–203.

Wargo PM. 1996. Consequences of environmental stress on oak: Predisposition to pathogens. Annales Des Sciences Forestieres 53(2-3): 359–368.

Zalkalns O, Celma L. 2021. The distribution of bacteria Gibbsiella quercinecans and Brenneria goodwinii in oak (Quercus robur L.) stands in Latvia. IOP Conference Series: Earth and Environmental Science

Zhang BC, Zhou XH, Zhou LY, Ju RT. 2015. A global synthesis of below-ground carbon responses to biotic disturbance: a meta-analysis. Global Ecology and Biogeography 24(2): 126–138.

Zinke PJ. 1962. The Pattern of Influence of Individual Forest Trees on Soil Properties. Ecology 43(1): 130–133.

