## Supplementary Information for "The cause-effect conundrum of local-scale site and soil factors in acute oak decline (AOD)"

*These authors made equal contributions.

Tree decline, acute oak decline, *Quercus robur*, *Quercus petraea*, soil biogeochemistry, waterlogging, plant-soil feedbacks

Table S1. Soil physicochemical properties at the three study sites. Data are overall mean (± SD) based on soil samples taken from a depth of 5-15 cm around 20 sampled trees (10 symptomatic, 10 non-symptomatic) per site. Means for each soil characteristic sharing a letter in common are not significantly different ($p<0.05$) among sites, according to Games-Howell Pairwise Comparisons.

| | Monks Wood (MW) | Stratfield Brake (SB) | Writtle Forest (WF) |
|---|---|---|---|
| Organic matter (%) | $13.5 \pm 2.7^{A}$ | $15.3 \pm 3.1^{A}$ | $16.3 \pm 6.4^{A}$ |
| pH ($H_2O$) | $4.7 \pm 0.7^{A}$ | $3.6 \pm 0.2^{B}$ | $3.6 \pm 0.3^{B}$ |
| Clay (%) | $1.6 \pm 0.8^{A}$ | $0.6 \pm 0.3^{B}$ | $0.4 \pm 0.5^{B}$ |
| Silt (%) | $68.7 \pm 7.8^{A}$ | $56.2 \pm 5.3^{B}$ | $47.6 \pm 10.1^{C}$ |
| Sand (%) | $30.0 \pm 7.9^{C}$ | $43.1 \pm 5.6^{B}$ | $52.0 \pm 10.7^{A}$ |
| Total C (%) | $6.0 \pm 1.5^{A}$ | $7.6 \pm 2.2^{A}$ | $8.3 \pm 4.0^{A}$ |
| Total N (%) | $0.47 \pm 0.1^{A}$ | $0.52 \pm 0.1^{A}$ | $0.42 \pm 0.2^{A}$ |
| Total mineral N (mg $kg^{-1}$) | $37.9 \pm 18.4^{A}$ | $27.8 \pm 14.7^{A}$ | $35.0 \pm 35.8^{A}$ |
| Olsen P (mg $kg^{-1}$) | $9.4 \pm 5.6^{B}$ | $20.9 \pm 12.5^{A}$ | $24.8 \pm 22.6^{A}$ |
| CEC ($mmol_c$ $kg^{-1}$) | $301.2 \pm 142.0^{A}$ | $195.8 \pm 27.2^{B}$ | $126.9 \pm 52.5^{C}$ |
| Exchangeable Al (mg $kg^{-1}$) | $190.2 \pm 174.2^{C}$ | $531.8 \pm 141.3^{A}$ | $306.7 \pm 116.6^{B}$ |
| Exchangeable Ca (mg $kg^{-1}$) | $3847.0 \pm 2512.0^{A}$ | $1628.8 \pm 389.7^{B}$ | $826.0 \pm 509.0^{C}$ |
| Exchangeable Fe (mg $kg^{-1}$) | $9.2 \pm 11.2^{B}$ | $36.1 \pm 24.3^{A}$ | $32.8 \pm 37.3^{A}$ |
| Exchangeable K (mg $kg^{-1}$) | $310.7 \pm 90.0^{A}$ | $289.4 \pm 69.0^{A}$ | $227.7 \pm 143.7^{A}$ |
| Exchangeable Mg (mg $kg^{-1}$) | $305.6 \pm 85.2^{A}$ | $172.0 \pm 34.5^{B}$ | $147.3 \pm 80.9^{B}$ |
| Exchangeable Mn (mg $kg^{-1}$) | $26.7 \pm 16.1^{B}$ | $21.8 \pm 8.9^{B}$ | $65.6 \pm 43.7^{A}$ |
| Exchangeable Na (mg $kg^{-1}$) | $59.9 \pm 19.5^{A}$ | $42.5 \pm 13.7^{B}$ | $24.1 \pm 14.0^{C}$ |
| Bulk density (g $cm^3$) | $0.803 \pm 0.102^{B}$ | $0.543 \pm 0.163^{C}$ | $1.041 \pm 0.299^{A}$ |

Table S2. Characteristics of the soils (40-50 cm depth) at the three study sites.

| | Monks Wood (MW) | Stratfield Brake (SB) | Writtle Forest (WF) |
|---|---|---|---|
| Organic matter (%)[a] | 11.54 (2.19)A | 11.94 (2.18)A | 7.71 (2.72)B |
| pH ($H_2O$)[a] | 4.89 (0.66)A | 3.68 (0.15)B | 3.63 (0.32)B |
| Clay (%)[a] | 2.24 (1.74)A | 0.93 (0.41)B | 0.54 (0.27)C |
| Silt (%)[a] | 72.09 (8.15)A | 61.01 (4.70)B | 57.44 (8.47)B |
| Sand (%)[a] | 23.91 (5.81)B | 37.98 (4.96)A | 41.50 (8.34)A |
| Total C (%) | 5.35 (1.20)A | 5.53 (1.40)A | 4.35 (2.10)A |
| Total N (%) | 0.41 (0.11)A | 0.39 (0.09)A | 0.22 (0.10)B |
| Total mineral N (mg $kg^{-1}$) | 30.34 (10.90)A | 18.27 (11.53)B | 13.79 (11.76)B |
| Olsen P (mg $kg^{-1}$) | 8.10 (6.63)A | 13.09 (5.60)A | 14.03 (17.13)A |
| CEC ($mmol_c$ $kg^{-1}$) | 299.8 (145.1)A | 169.3 (26.99)B | 86.52 (34.3)C |
| Exchangeable Al (mg $kg^{-1}$) | 184.1 (170.5)C | 524.9 (126.4)A | 366.6 (137.8)B |
| Exchangeable Ca (mg $kg^{-1}$) | 3870 (2494)A | 1326.8 (320.0)B | 388.6 (331.0)C |
| Exchangeable Fe (mg $kg^{-1}$) | 5.63 (4.22)B | 22.60 (18.21)A | 26.47 (22.06)A |
| Exchangeable K (mg $kg^{-1}$) | 272.5 (77.9)A | 227.7 (56.8)A | 158.7 (99.4)B |
| Exchangeable Mg (mg $kg^{-1}$) | 310.1 (94.6)A | 159.32 (38.00)B | 83.4 (58.7)C |
| Exchangeable Mn (mg $kg^{-1}$) | 22.23 (14.33)AB | 14.72 (7.38)B | 28.83 (18.22)A |
| Exchangeable Na (mg $kg^{-1}$) | 64.36 (23.21)A | 41.68 (19.49)B | 12.25 (8.39)C |

[a]Overall mean (± SD) based on soil samples taken from a depth of 40-50 cm around 20 sampled trees (10 symptomatic, 10 non-symptomatic) per site. Means for each soil characteristic sharing a letter in common are not significantly different ($p<0.05$) among sites, according to Games-Howell Pairwise Comparisons.

Table S3. Decimal Latitude/longitude coordinates, AOD symptom status (Symptomatic (S) or Asymptomatic (A)) and species classification for the selected trees from the three study sites (Writtle (WF), Monkswood (MW) and Stratfield Brake (SB).

| Site | Tree ID | Latitude | Longitude | AOD symp-tom status | Classification of species according to morphometric analysis of five leaves tree$^{-1}$ | | DNA-based classification | |
|---|---|---|---|---|---|---|---|---|
| | | | | | # of leaves classified as *Q. robur* | # of leaves classified as *Q. petraea* | Species | Haplo-type |
| WF | WD01191 | 51.701480 | 0.374720 | S | 4 | 1 | robur | newA |
| WF | WD01192 | 51.704690 | 0.399020 | S | 5 | 0 | robur | 12 |
| WF | WD01193 | 51.709850 | 0.398250 | S | 5 | 0 | robur | 12 |
| WF | WD01194 | 51.710350 | 0.403590 | S | 5 | 0 | robur | newB |
| WF | WD01195 | 51.689710 | 0.368150 | S | 0 | 5 | robur | 10bis |
| WF | WD01196 | 51.690590 | 0.361910 | S | 1 | 4 | petraea | newB |
| WF | WD01197 | 51.692080 | 0.358150 | S | 0 | 5 | petraea | 12 |
| WF | WD01198 | 51.694750 | 0.353060 | S | 5 | 0 | robur | 10bis |
| WF | WD01199 | 51.693130 | 0.357240 | S | 0 | 5 | petraea | 12 |
| WF | WD01187 | 51.692290 | 0.364120 | S | 0 | 5 | petraea | 12 |
| WF | WH01200 | 51.697990 | 0.383530 | A | 4 | 1 | robur | newA |
| WF | WH01872 | 51.700870 | 0.392790 | A | 5 | 0 | robur | newB |
| WF | WH01874 | 51.706930 | 0.395550 | A | 5 | 0 | robur | newB |
| WF | WH01875 | 51.703250 | 0.394730 | A | 1 | 4 | robur | 12 |
| WF | WH01876 | 51.701050 | 0.372850 | A | nd | nd | nd | nd |
| WF | WH01879 | 51.698550 | 0.348340 | A | 5 | 0 | robur | 12 |
| WF | WH01183 | 51.698390 | 0.375840 | A | 5 | 0 | robur | 10bis |
| WF | WH01184 | 51.700970 | 0.351650 | A | 5 | 0 | robur | newB |
| WF | WH01185 | 51.696990 | 0.352230 | A | 5 | 0 | robur | newA |
| WF | WH01188 | 51.693490 | 0.353190 | A | 0 | 5 | petraea | 12 |
| MW | MD00991 | 52.408494 | -0.235161 | S | 5 | 0 | nd | nd |
| MW | MD00992 | 52.408686 | -0.234227 | S | 4 | 1 | robur | 12 |
| MW | MD00993 | 52.409061 | -0.233432 | S | 5 | 0 | robur | 10 |
| MW | MD00994 | 52.409570 | -0.231060 | S | 5 | 0 | robur | 10 |
| MW | MD00995 | 52.409990 | -0.230540 | S | 5 | 0 | robur | 10 |
| MW | MD00996 | 52.412012 | -0.235358 | S | 3 | 2 | robur | 10 |
| MW | MD00997 | 52.411590 | -0.235380 | S | 4 | 1 | robur | 10 |
| MW | MD00998 | 52.411260 | -0.235130 | S | 4 | 1 | robur | 10 |
| MW | MD00999 | 52.410780 | -0.236940 | S | 5 | 0 | robur | 10 |
| MW | MD01000 | 52.403462 | -0.228188 | S | 5 | 0 | robur | 10 |

| | | | | | | | | |
|---|---|---|---|---|---|---|---|---|
| MW | MH00812 | 52.403300 | -0.232050 | A | 3 | 2 | robur | 11 |
| MW | MH00891 | 52.403830 | -0.232320 | A | 5 | 0 | robur | 10 |
| MW | MH00892 | 52.403610 | -0.231030 | A | 4 | 1 | robur | 10bis |
| MW | MH00893 | 52.402910 | -0.228030 | A | 5 | 0 | robur | 11 |
| MW | MH00894 | 52.403670 | -0.230160 | A | 5 | 0 | robur | 12bis |
| MW | MH00895 | 52.411940 | -0.234680 | A | 5 | 0 | robur | 10 |
| MW | MH00896 | 52.411460 | -0.232840 | A | 5 | 0 | robur | 10 |
| MW | MH00897 | 52.409720 | -0.229590 | A | 4 | 1 | robur | 12 |
| MW | MH00898 | 52.409180 | -0.232640 | A | 5 | 0 | robur | 10 |
| MW | MH00899 | 52.408650 | -0.233870 | A | 5 | 0 | robur | 10 |
| SB | SD0054 | 51.803420 | -1.281100 | S | 5 | 0 | robur | 12 |
| SB | SD0056 | 51.803100 | -1.283110 | S | 5 | 0 | robur | 10bis |
| SB | SD00736 | 51.803450 | -1.282270 | S | 5 | 0 | robur | 12 |
| SB | SD00737 | 51.803060 | -1.283840 | S | 5 | 0 | robur | 12 |
| SB | SD00738 | 51.803600 | -1.281750 | S | 5 | 0 | robur | 10bis |
| SB | SD00739 | 51.803100 | -1.281700 | S | 5 | 0 | robur | 12 |
| SB | SD00813 | 51.802810 | -1.284490 | S | 5 | 0 | robur | 12 |
| SB | SD00816 | 51.803090 | -1.282370 | S | 4 | 1 | robur | 12 |
| SB | SD00970 | 51.802860 | -1.282660 | S | 5 | 0 | robur | 12 |
| SB | SDX | 51.803470 | -1.281380 | S | 5 | 0 | robur | 12 |
| SB | SH01 | 51.803380 | -1.283220 | A | 5 | 0 | robur | 12 |
| SB | SH02 | 51.803380 | -1.281780 | A | 5 | 0 | robur | 12 |
| SB | SH03 | 51.804370 | -1.280930 | A | 5 | 0 | robur | 12 |
| SB | SH04 | 51.804160 | -1.281110 | A | 5 | 0 | robur | 12 |
| SB | SH05 | 51.803330 | -1.282970 | A | 5 | 0 | robur | 12 |
| SB | SH06 | 51.803280 | -1.283540 | A | 5 | 0 | robur | 12 |
| SB | SH07 | 51.802980 | -1.284770 | A | 5 | 0 | robur | 12 |
| SB | SH08 | 51.803230 | -1.283900 | A | 5 | 0 | robur | 12 |
| SB | SH09 | 51.804100 | -1.280680 | A | 5 | 0 | robur | 12 |
| SB | SH10 | 51.803690 | -1.281180 | A | 5 | 0 | robur | 12 |

Table S4: Prediction performance across 100 GBM runs

| | AUC | SD | Specificity | SD | Sensitivity | SD |
|---|---|---|---|---|---|---|
| Mean | 0.88 | 0.06 | 0.92 | 0.09 | 0.82 | 0.11 |
| Median | 0.89 | - | 0.89 | - | 0.83 | - |

Table S5: Variables with greatest relative influence on tree health status

| **Variable** | **% times selected** |
|---|---|
| Depth to gleying | 100 |
| Exchangeable Fe (40-50 cm) | 95 |
| Olsen P (40-50 cm) | 90 |
| Basal area (0-20 m) | 88 |
| Available ammonium N (40-50 cm) | 83 |
| Crown width (East-West) | 79 |
| Basal area (20-40 m) | 77 |
| Total N (0-10 cm) | 71 |
| Total C (40-50 cm) | 69 |
| Total C (0-10 cm) | 65 |
| Exchangeable Mn (40-50 cm) | 65 |
| Exchangeable Fe (0-10 cm) | 63 |
| Tree Height | 59 |
| Exchangeable Na (40-50 cm) | 54 |
| Exchangeable K (0-10 cm) | 52 |
| Ammonium to nitrate ratio (40-50 cm) | 52 |
| Olsen P (0-10 cm) | 49 |
| Diameter at 1.35 m | 47 |
| Loss on ignition (40-50 cm) | 45 |
| Available ammonium N (0-10 cm) | 45 |

Table S6: Anova table for binomial regression model with tree health status as the outcome variable and depth to gleying and Exchangeable Fe (40-50 cm) as predictors.

| Variable | LR Chisq | Df | Pr(>Chisq) |
|---|---|---|---|
| Depth to gleying | 37.8 | 1 | 7.89e-10 |
| Exchangeable Fe (40-50 cm) | 5.28 | 1 | 0.0216 |

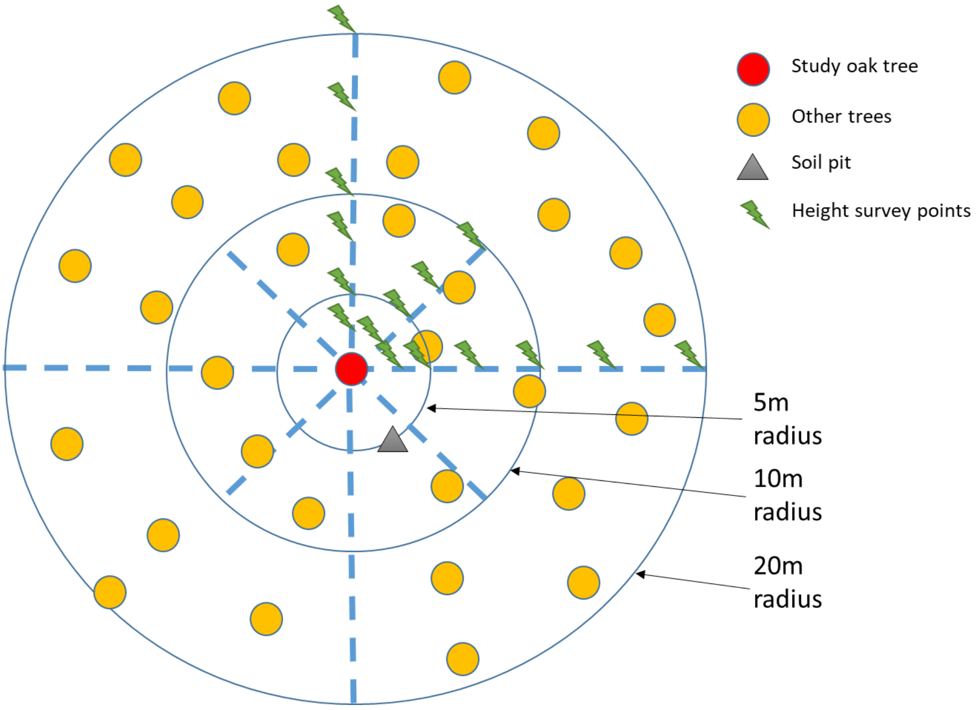


Figure S1. Schematic diagram of the sample measurements made of the site around each study tree

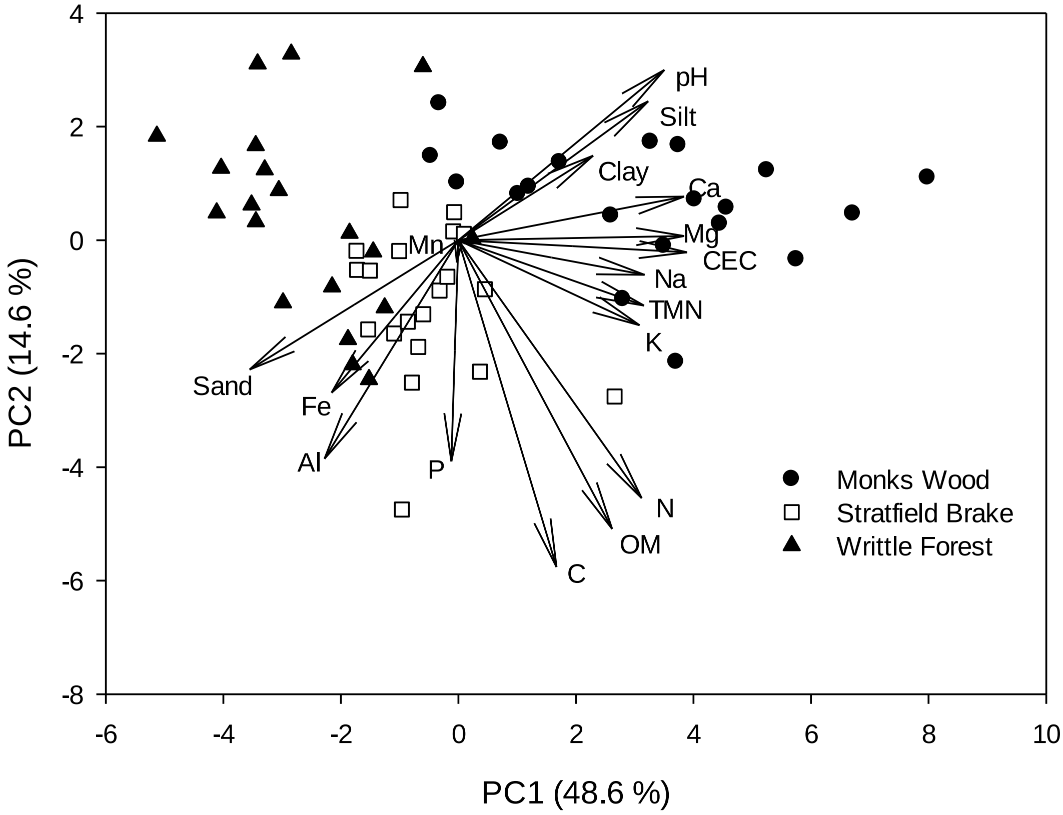


Figure S2. Principal Components Analysis of the study sites (symbols) and soil parameters (vectors) for deep (40-50 cm) soils. One-Way ANOVA on PC1 scores and Games-Howell pairwise comparisons revealed that all three sites differed significantly ($P<0.05$) from one another.

| Degrees of Freedom | Site-Depth (S-D) 3 | Health (H) 1 | S-D*H 3 |
| --- | --- | --- | --- |
| Total P | **<0.001** | **0.021** | 0.506 |

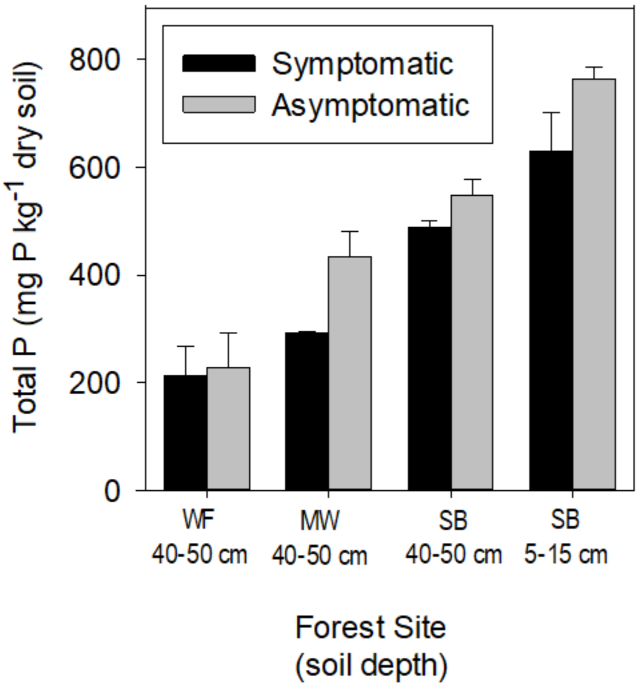


Figure S3. Concentrations of Total P (mean ± SE) of soil sampled from 40-50 cm depth (all forest sites) or 5-15 cm depth (Stratfield Brake site only) for selected symptomatic or asymptomatic trees from each of the study sites: Monks Wood (MW), Writtle Forest (WF), Stratfield Brake (SB). For each site and soil depth an equal number of symptomatic and asymptomatic trees were compared with sample size for each health class (n) as follows: $WF_{40\text{-}50\ cm}$ (n=5), $MW_{40\text{-}50\ cm}$ (n=3), $SB_{40\text{-}50\ cm}$ (n=6), $SB_{5\text{-}15\ cm}$ (n=3). The Table above the figure shows the outcome of two-way ANOVA (Box-Cox-transformed data) examining the association of Site-Depth, Health status and their interaction on total soil P concentration. Total P concentrations were determined via Aqua Regia (3.5 ml conc. HCl and 1.2 ml conc. $HNO_3$) digestion (140 °C for 2.5 hours) of 0.5 g air-dried soil samples (<2 mm) followed by analysis of digests using ICP-OES.
